# Maternal periconceptional nutrition is associated with the vaginal microbiome along pregnancy

**DOI:** 10.64898/2026.11.25.769327

**Authors:** Jinhui Liu, Andrew Batuure, Julia Urban, Emily Watters, Heekuk Park, Matthew K. Hoffman, Aya Brown Kav, Anne-Catrin Uhlemann, Tal Korem

## Abstract

The human vaginal microbiome has been repeatedly associated with adverse pregnancy outcomes, and therefore understanding lifestyle factors associated with it may provide insights into strategies for improving pregnancy and reproductive health. The periconceptional period is a critical time window in pregnancy, with nutrition seen as a key factor for pregnancy health. However, the associations between periconceptional nutrition and the vaginal microbiome along pregnancy remain underexplored. Using 280 vaginal microbiome samples from 101 women collected longitudinally throughout pregnancy, along with paired food frequency questionnaires targeting the periconceptional period, we characterized the associations between periconceptional diet, dietary supplementation, and the vaginal microbiome. We found that periconceptional dietary indices are not strongly associated with the vaginal microbiome during pregnancy. However, several dietary components are significantly associated with both *Lactobacillus crispatus* and *Lactobacillus iners* in the first trimester, and in opposite directions. Additionally, we identified several associations between consumption of periconceptional dietary supplements and the vaginal microbiome in the third trimester. Interestingly, an increase in taxa typically associated with vaginal dysbiosis, such as *Sneathia* and *Megaphera*, was associated with the intake of specific micronutrients. Among these micronutrients, copper and zinc were notable, as individuals reporting their supplementation had average intake levels that exceeded the recommended daily allowance. Overall, our findings reveal pregnancy-stage-specific associations of the human vaginal microbiome with periconceptional nutrition, and suggest the potential of specific nutrients to modulate the vaginal microbiome.

## Introduction

The vaginal microbiome plays a crucial role in female reproductive health. During pregnancy, this microbial community typically shifts towards higher rates of *Lactobacillus* dominance, which is considered protective against infection and inflammation^1–4^. Similarly, dysbiosis of the vaginal microbiome has been associated with adverse pregnancy outcomes (APOs), such as preterm birth and preeclampsia^5–12^. Socioeconomic and lifestyle factors, such as employment, housing, and hygiene practices have also been associated with the vaginal microbiome^13–15^. Characterizing these and similar associations may help identify modifiable risk factors for pregnancy complications.

Diet is a major modifiable risk factor for APOs^16,17^. One of its important aspects is maternal nutrition during the periconceptional period, a critical window which may influence oocyte development, fertilization, implantation, and early placental development^18^. Both undernutrition and overnutrition during this period are associated with adverse pregnancy and birth outcomes^19–21^. Recent studies suggest that a Mediterranean-style diet rich in vegetables, fish, legumes, and cereals is effective in preventing APOs, potentially because its antioxidant properties could alleviate elevated oxidative stress during placental development^22,23^. Other, simpler dietary indices that emphasize legumes, vegetables and whole grains have similarly shown associations with lower rates of APOs^16^.

Diet has also been associated with the vaginal microbiome: high-fiber, low-glycemic-index, and low-fat diets were each associated with lower risk of bacterial vaginosis^24,25^; and in pregnancy, increasing evidence suggests that diet is linked to the vaginal microbiome^26–30^. However, these findings are often limited by the granularity at which dietary and vaginal microbiome data were analyzed^31^. This, in turn, limits the specificity of the associations identified, and, consequently, downstream interventions that could be investigated. Finally, the associations of diet in the periconceptional period with the vaginal microbiome during pregnancy remain to be characterized.

Dietary supplements are commonly taken before and during pregnancy. Several studies have identified that deficiency in vitamins such as A, D, and E during pregnancy is associated with risk of bacterial vaginosis^32–37^. A randomized clinical trial also found that vitamin A supplementation reduced the incidence of bacterial vaginosis^38^, suggesting that supplements may be candidates for microbiome-targeting interventions. Although dietary supplements are widely used in the periconceptional period^39^, their associations with the vaginal microbiome remain underexplored, hindering the development of targeted nutritional interventions to support vaginal and reproductive health.

Here, we investigated periconceptional nutrition and supplement intake and their association with the vaginal microbiome during pregnancy in 101 individuals. Analyzing longitudinal microbiome and high-resolution dietary profiles, we found that periconceptional diet was associated with the first-trimester vaginal microbiome. We identified several dietary components that were associated with *L. crispatus* and *L. iners*, the predominant *Lactobacillus* species in the vagina, in opposite directions. We further found that high micronutrient intake, largely driven by dietary supplementation, was associated with the presence of dysbiosis-associated taxa in the third trimester. Overall, our results reveal multiple associations between periconceptional diet and supplements and the vaginal microbiome throughout pregnancy.

## Results

### Study design and cohort statistics

As part of a pilot study for studying APOs in pregnancy, we selected 124 participants from the Nulliparous Pregnancy Outcomes Study: Monitoring Mothers-to-be (nuMoM2b) cohort^40^. Up to three vaginal swabs were available per participant, self-collected at three nuMoM2b study visits that took place between 6^0^-13^6^ (visit 1), 16^0^-21^6^ (visit 2), and 22^0^-29^6^ (visit 3) weeks^days^ of gestation. We performed 16S rRNA gene amplicon sequencing on those swabs to profile the vaginal microbiome, and removed 18 samples with either processing errors^41^ or low read counts (**Methods**; **Fig. S1**). Dietary intake during the three months before pregnancy was also assessed at visit 1 using a modified Block 2005 food frequency questionnaire (FFQ)^42^. We removed 54 samples from 21 individuals with FFQ data that were missing or reflected abnormally high or low total energy intake (**Methods**). We used the remaining 280 samples for downstream analysis (**Fig. S1**), corresponding to 101 participants with mean±sd age of 26.45±5.57 (**Table S1**).^42^Alternate Mediterranean Diet (aMED), the Healthy Eating Index-2010 (HEI-2010), and the Alternative Healthy Eating Index 2010 (AHEI-2010)^42–45^. As participants were from a pilot study that included randomly selected participants from several case-control studies, the incidence rates of APOs in this cohort are higher than in the general population, with 53 participants developing at least one APO, including 19 and 20 cases of spontaneous preterm birth and preeclampsia, respectively (**Table S1**).

### Periconceptional dietary indices are associated with the presence of a few vaginal taxa

To investigate the associations between overall dietary patterns and the vaginal microbiome during pregnancy, we evaluated several dietary indices, including the aMED, HEI-2010, and AHEI-2010 scores^43–45^. To first examine how the overall microbiome composition is related to these dietary indices, we evaluated beta diversity metrics. We did not identify significant associations at any visit, evaluating both Bray-Curtis dissimilarity and Jaccard distances of the microbiome (PERMANOVA *p* > 0.043, FDR > 0.1 for all; **Fig. S3**). We then tested if the relative abundance or presence of any particular Amplicon Sequence Variant (ASV) in the microbiome is associated with these dietary indices. Although the relative abundances of none of the ASVs were significantly associated with dietary indices (ALDEx3 *p* > 0.12, FDR > 0.1 for all; **Fig. S4a**), we found that the presence of ASV65 (*Peptoniphilus harei*) and ASV38 (*Finegoldia magna H*) at visit 1 were positively associated with HEI-2010 score (binomial regression *p* = 0.0072 and 0.017, respectively, *q* = 0.050 and 0.061; **Figs. 2a,b**, **S5a**). To capture less abundant taxa, we also examined potential associations with diet indices at the genus level. Despite again finding no associations between genus-level relative abundance and diet indices at all visits (ALDEx3 *p* > 0.056, FDR > 0.1; **Fig. S4b**), we found that their presence and absence was associated with dietary indices.

**Figure 1.**
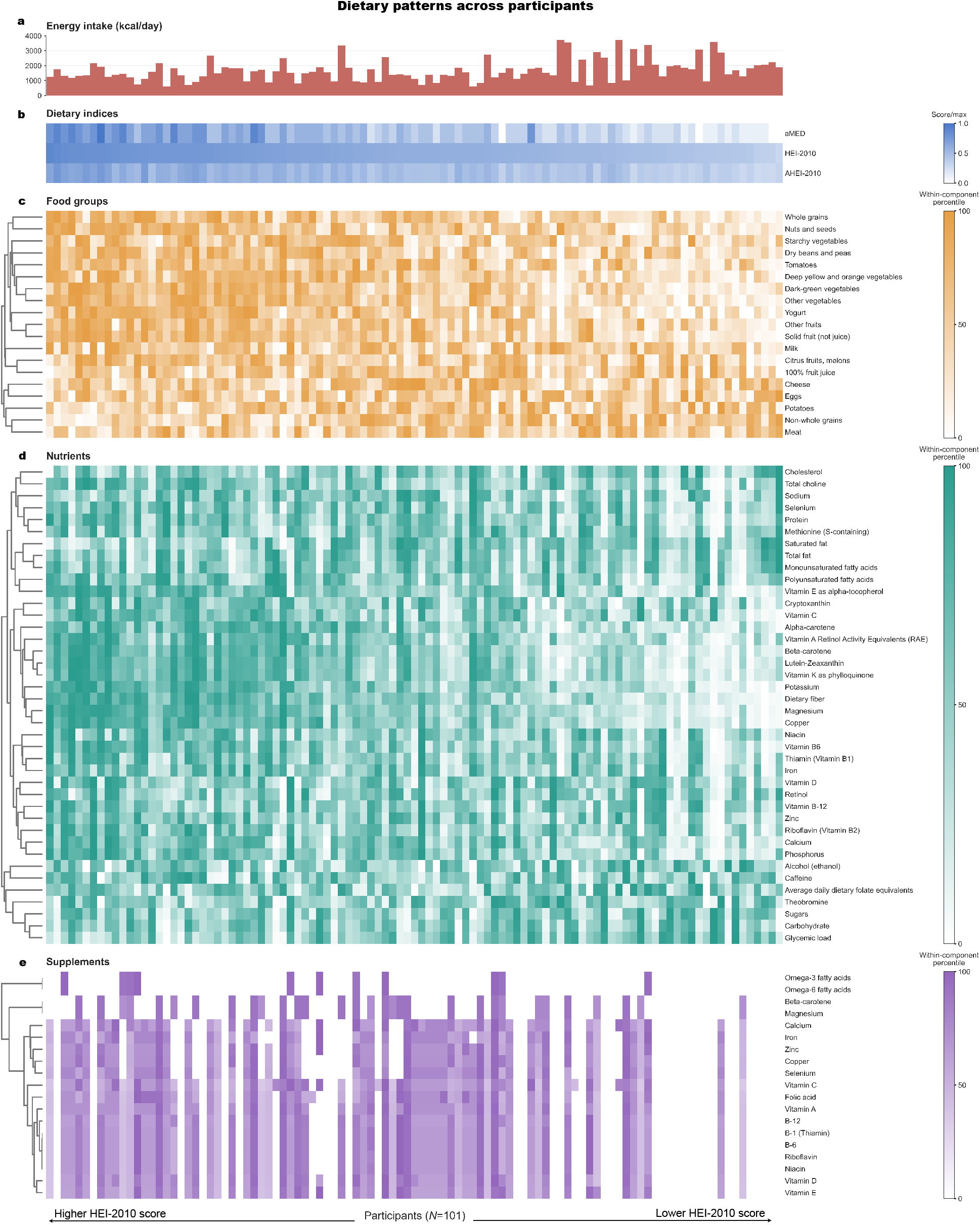
Dietary patterns across study participants. **a.** Bar plot showing the daily calorie intake of each participant in this study. **b-e.** Heatmaps showing the dietary intake patterns of participants in this study, including dietary indices (**b**), food groups (**c**), nutrients (**d**), and supplements (**e**). For each category, components (rows) are hierarchically clustered by Euclidean distances using average linkage. Participants (columns) are ordered from high to low HEI-2010 score.

**Figure 2.**
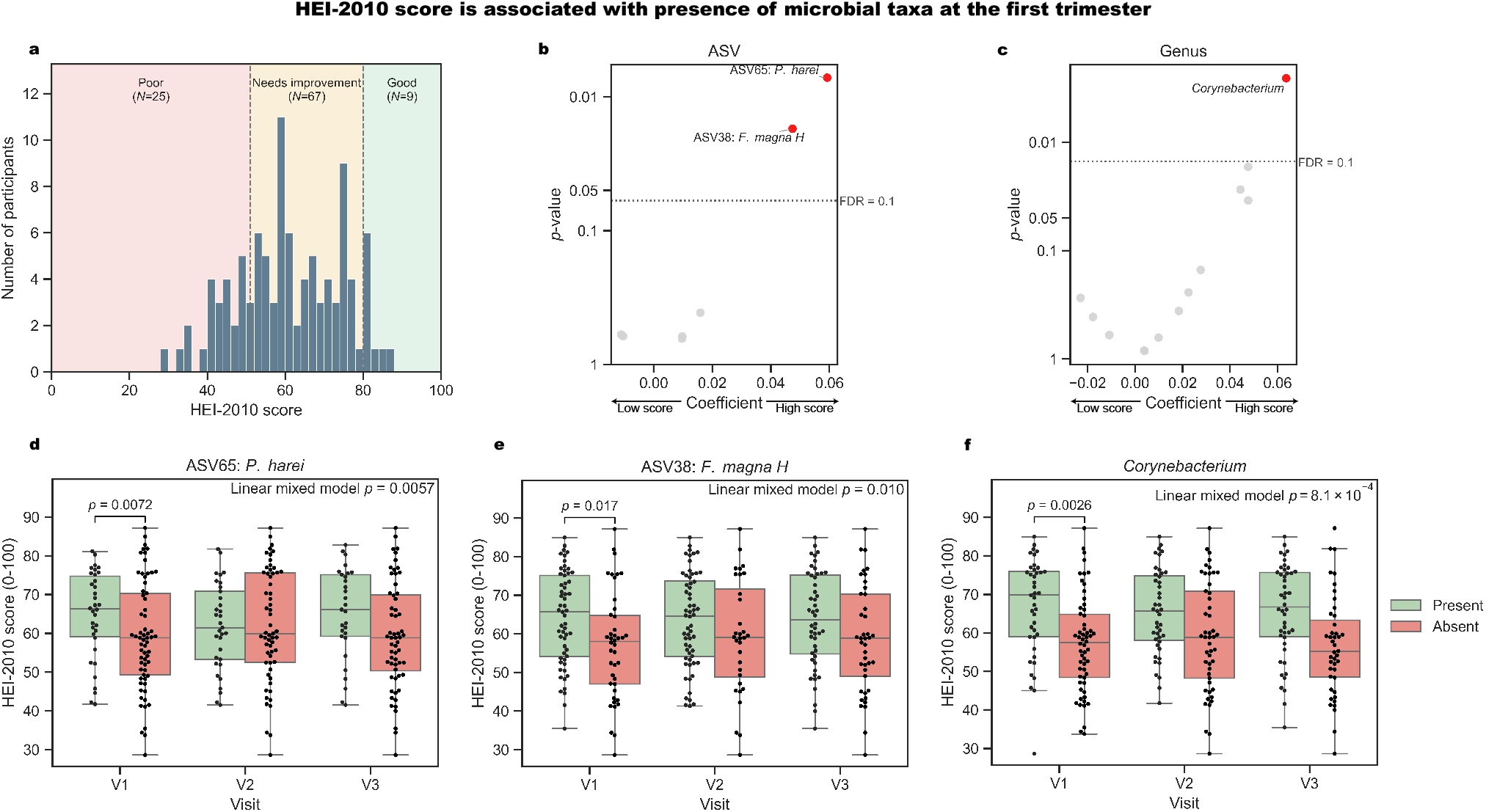
HEI-2010 is associated with the first trimester vaginal microbiome. **a.** Histogram showing distribution of HEI-2010 score of participants in this study. **b,c.** Volcano plots showing associations (binomial regression coefficient, x-axis) and statistical significance (y-axis) of HEI-2010 score with the presence of microbial ASVs (**b**) and genera (**c**) at visit 1. **d-f.** Box and swarm plots (box, IQR; line, median; whiskers, 1.5xIQR) showing HEI-2010 score grouped by presence of ASV65 (**d**), ASV38 (**e**), and *Corynebacterium* (**f**) across visits. The visit-specific *p*-value is from the corresponding binomial regression model in panels **a** and **b**, while the overall *p*-value of each panel corresponds to a linear mixed model fitted on samples across all visits (FDR < 0.1). Presence and absence are called according to a randomly selected rarefaction run for visualization purposes (**Methods**).

Specifically, at visit 1, *Corynebacterium* was enriched in individuals with high HEI-2010 scores (binomial regression *p* = 0.0026, *q* = 0.033; **Figs. 2c**, **S5b**), and *Staphylococcus* and *Corynebacterium* were enriched in individuals with high AHEI-2010 scores (*p* = 0.0060 and 0.0075, respectively, *q* = 0.049 and 0.049; **Fig. S5b**). In addition to analyzing samples from each visit separately, we further combined samples from all three visits to test these associations using a linear mixed model with additional adjustment for gestational age, and found consistent findings (*p* = 0.0057 and 0.010 for the association of ASV65 and ASV38 with HEI-2010, respectively, *q* = 0.046 and *q* = 0.046; *p* = 8.1×10^−4^, *q* = 0.011 for association of *Corynebacterium* with HEI-2010; and *p* = 0.0017 and 0.0034 for association of *Corynebacterium* and *Staphylococcus* with AHEI-2010, respectively, *q* = 0.022 and *q* = 0.022; **Figs. 2d-f, S5c,d**). Overall, while community-wide associations with periconceptional dietary indices were weak, we found limited associations between some indices and the presence of specific taxa.

### Dietary nutrient and food components are associated with the first trimester vaginal microbiome

Identifying specific dietary nutrients or food components associated with the vaginal microbiome could provide insights into the relationship between nutrition and the vaginal ecosystem. Evaluating associations with all 59 specific food items and nutrients profiled via FFQ (**Table S2**), we found that consumption of 100% fruit juice (PERMANOVA of Bray-Curtis dissimilarities *p* = 0.0029, *q* = 0.093), niacin (*p* = 0.0038, *q* = 0.093), and tomatoes (*p* = 0.0048, *q* = 0.093) were significantly associated with the vaginal microbiome at the first trimester (**Fig. 3a**). Similar to dietary indices, no dietary component was significantly associated with the vaginal microbiome at later visits (*p* > 0.0065, FDR > 0.1 for all; **Fig. S6**).

**Figure 3.**
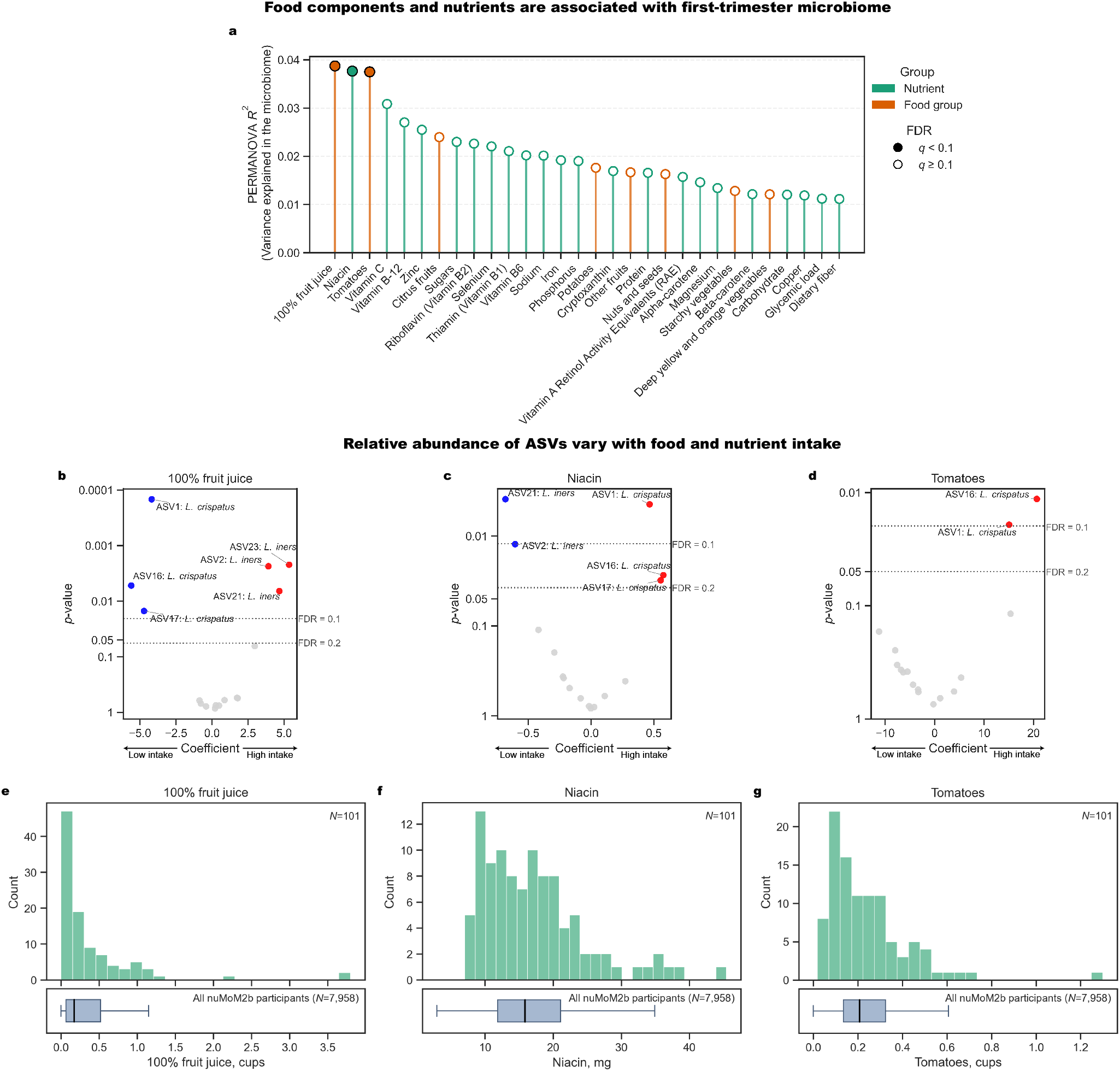
Food and nutrients are significantly associated with the first trimester vaginal microbiome and the abundance of specific key taxa. **a**. Lollipop plot of PERMANOVA *R*^2^ between diet variables (top 30 based on *R*^2^) and Bray-Curtis dissimilarity between visit 1 samples. **b-d.** Volcano plot showing associations between abundance of microbial ASVs at visit 1 and consumption of 100% juices (**b**), niacin (**c**), and tomatoes (**d**). **e-g.** Distribution of 100% fruit juices (**e**), niacin (**f**), and tomatoes (**g**). Histograms in upper panels show distributions in this study, while boxplots in lower panels show distributions in the full nuMoM2b cohort.

We next analyzed differential abundance of ASVs associated with the three identified dietary components. We found three ASVs which were positively correlated with juice consumption, all annotated as *L. iners* (ALDEx3; ASV2: *p* = 0.0022, *q* = 0.014; ASV21: *p* = 0.0058, *q* = 0.029; ASV23: *p* = 0.0019, *q* = 0.012), and three which were negatively associated, all annotated as *L. crispatus* (ASV1: *p* = 8.4×10^−5^, *q* = 0.0012; ASV16: *p* = 0.0048, *q* = 0.031; ASV17: *p* = 0.014, *q* = 0.079; **Fig. 3b**). We observed the opposite trends for niacin: ASV1 (*L. crispatus*) was positively associated with niacin (*p* = 0.0043, *q* = 0.056), while ASV21 (*L. iners*) had a negative association (*p* = 0.0040, *q* = 0.043; **Fig. 3c**). We found similar associations with tomatoes: two ASVs annotated as *L. crispatus* (ASV1 and ASV16) were positively associated with its consumption (*p* = 0.017 and *p* = 0.010, respectively, *q* = 0.099 and 0.089; **Fig. 3d**). We found weaker but consistent trends for all three dietary components when leveraging samples from all three visits using a linear mixed model (fruit juice: *p* = 0.0051, *q* = 0.087 for ASV21; tomatoes: *p* = 0.0047, *q* = 0.045 for ASV1, *p* = 0.0052, *q* = 0.044 for ASV16, *p* = 0.021, *q* = 0.090 for ASV17, and *p* = 0.013, *q* = 0.071 for ASV21; **Fig. S7a-c**). However, at the genus level, we did not observe any significant associations between these dietary components and vaginal microbes (FDR > 0.1; **Fig. S8a-c**). Overall, these results indicate that periconceptional dietary components are associated with the vaginal microbial composition at the ASV rather than genus level during early pregnancy.

### Periconceptional dietary supplements are associated with vaginal microbiome during late pregnancy

We next sought to assess the association of dietary supplements, a highly prevalent and readily modifiable source of micronutrient intake, with the vaginal microbiome. Of participants included in this analysis, 37% reported taking supplements since pre-pregnancy, and an additional 55% started taking prenatal vitamins for prevention before visit 1 (**Table S1**). As some micronutrients from supplements had nearly perfect correlation with one another (both within this study and to a large degree within the broader nuMoM2b cohort), we first grouped those with Pearson *r* > 0.95 (**Fig. S9**). In contrast to the associations observed for dietary components, we did not find any supplement that was significantly associated with the beta diversity of the vaginal microbiome in the first or second trimesters, with and without adjustment for overall diet quality using HEI-2010 score (PERMANOVA of Bray-Curtis dissimilarities and Jaccard distances; FDR > 0.1 for all; **Fig. S10**). Surprisingly, we found that two and seven supplement groups were significantly associated with the vaginal microbiome in the third trimester when analyzing beta diversity via Bray-Curtis dissimilarities and Jaccard distances, respectively (PERMANOVA *p* < 0.05, FDR < 0.1; **Figs. 4a, S10**). This suggests an association between dietary supplementation and the presence of microbes in the vagina late in pregnancy.

**Figure 4.**
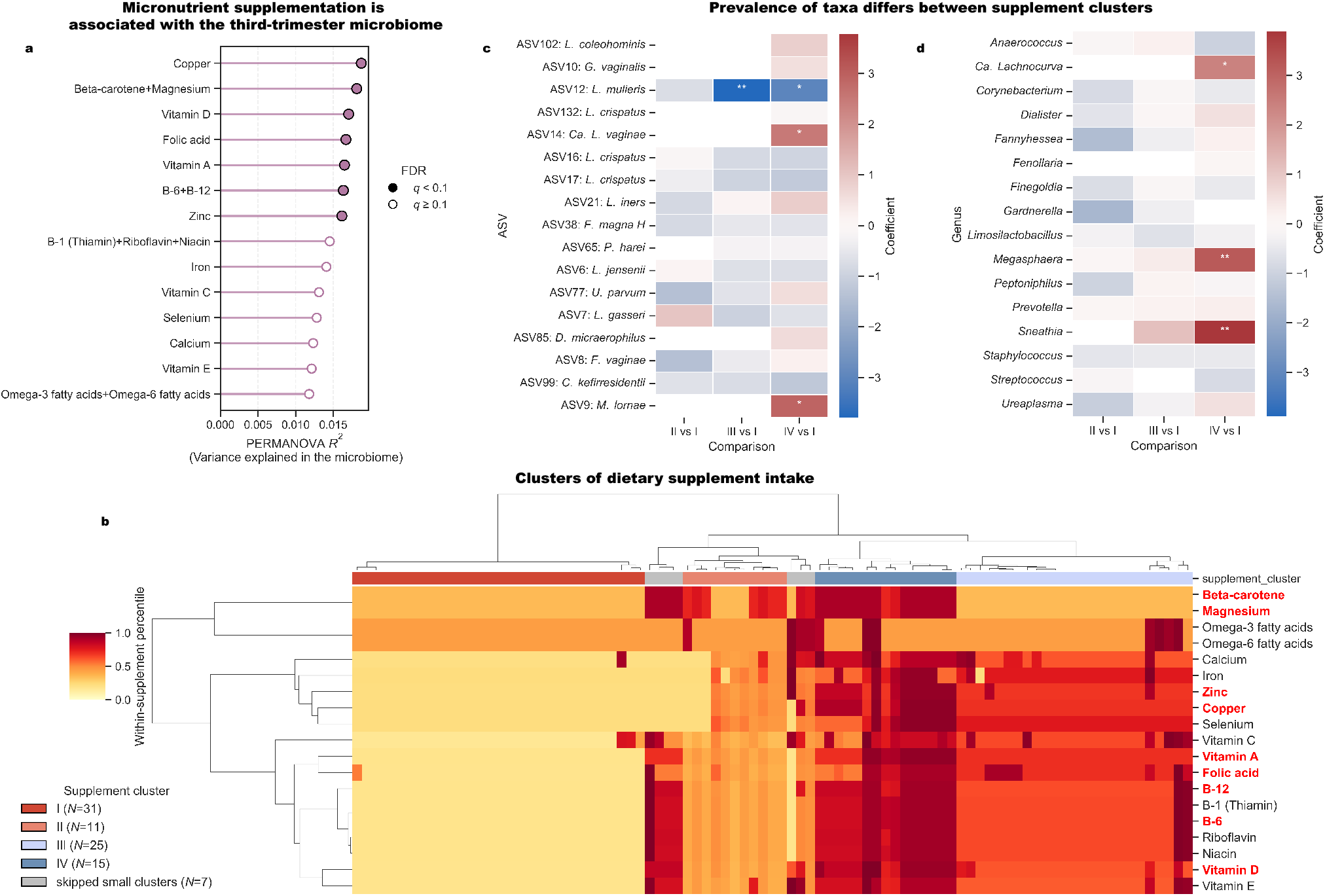
Supplement intake is associated with the vaginal microbiome in the third trimester. **a**. Lollipop plot of PERMANOVA *R*^2^ between supplement intake and Jaccard distance for visit 3 samples. **b.** Heatmap showing standardized supplement intake levels (rows) for each individual (columns). Supplements significantly associated with the vaginal microbiome in panel **a** are colored in red. **c,d.** Heatmaps showing comparison between ASV (**c**) and genus (**d**) prevalence among different clusters (***: *p* < 0.001, **: *p* < 0.01, *: *p* < 0.05; FDR < 0.1).

Given our limited sample size, instead of testing the associations between specific taxa and each micronutrient, we first performed hierarchical clustering on standardized dietary supplement profiles. We found four main ‘supplement clusters’ (**Methods**; **Figs. 4b**, **S11**; **Table S3**), which we named clusters I-IV based on the total relative supplement intake (**Fig. 4b; Table S3**): cluster I corresponded to little or no supplement intake (and was therefore used as baseline), cluster II exhibited low intake for most supplements except beta-carotene and magnesium, whereas clusters III and IV corresponded to moderate and high supplement intake, respectively. We found that these clusters were associated with maternal age at visit 1 and educational attainment (Kruskal-Wallis *p* = 0.010, *q* = 0.051; Fisher’s exact *p* = 0.025, *q* = 0.062; respectively), an association driven primarily by cluster I, which was enriched in individuals who were younger and had lower level of education (**Fig. S12**), two factors that are also associated with each other (two-sided Mann-Whitney *p* = 1.4×10^−5^).

As we observed significant associations between supplement intake and the vaginal microbiome primarily using Jaccard distances, we next prioritized investigating the associations between supplement clusters and the presence of specific microbes in the third trimester. We found that, compared to cluster I, the presence of ASV12 (*Lactobacillus mulieris*) was lower in clusters III and IV (binomial regression *p* = 0.0026 and 0.012, respectively, *q* = 0.029 and 0.081), and the presence of ASV9 (*Megasphaera lornae*) and ASV14 (*Ca. Lachnocurva vaginae*) were higher in cluster IV (*p* = 0.014 and 0.011, respectively, *q* = 0.081 and 0.081; **Fig. 4c**). To capture less abundant taxa, we also evaluated potential associations on the genus level, and found that the presence of *Sneathia* was also higher in cluster IV (*p* = 0.0069, *q* = 0.051; **Fig. 4d**).

### Use of periconceptional dietary supplements is associated with excess micronutrient intake

As consumption could be contributed by both diet and supplements, we sought to calculate the total intake of each micronutrient and evaluate the associations between total intake and the microbiome. We found that the presence of *Ca. L. vaginae* (ASV14) was associated with the intake of copper and vitamin D (binomial regression *p* = 0.0023 and 0.0070, respectively, *q* = 0.023 and 0.069; **Fig. 5a,b**). At the genus level, we found that *Megasphaera*, *Sneathia*, and *Gardnerella* were associated with copper, vitamin D, iron, beta-carotene, and zinc, consistent with our findings considering only intake from supplements (binomial regression *p* < 0.05; FDR < 0.1; **Fig. 5c**). Among those micronutrients, copper, zinc, and vitamin D were significantly associated with the third-trimester microbiome in both this analysis and when not considering dietary contribution (**Fig. 4a**), so we investigated the contribution of dietary supplements to their total intake. For all three micronutrients, we observed that total intake was on average more than two times higher in individuals who took supplements containing them than those who did not (two-sided Mann-Whitney *p* = 3.4×10^−11^, 6.0×10^−11^, and 3.9×10^−11^ for copper, zinc, and vitamin D, respectively; **Table 1**). Notably, we found that the average daily intake of copper and zinc was 2.4±1.0 mg/day and 21.9±10.1 mg/day for individuals who took dietary supplements, approximately twice the Recommended Dietary Allowances (RDA) for pregnancy^46^ (while still lower than the Tolerable Upper Intake Level [UL]), whereas the average intake of both elements was close to RDA for individuals who do not take supplements (1.2±0.4 mg/day and 9.7±4.3 mg/day, respectively), indicating high intake from supplements (**Table 1**). In contrast, we found that the intake of vitamin D in both groups was below RDA (**Table 1**), consistent with a previous analysis^39^. Overall, these results suggest that high intake of trace metals from dietary supplements is associated with higher vaginal prevalence of dysbiosis-associated taxa late in pregnancy, such as *Ca. L. vaginae*, *Megashphaera*, and *Sneathia*^47–51^.

**Figure 5.**
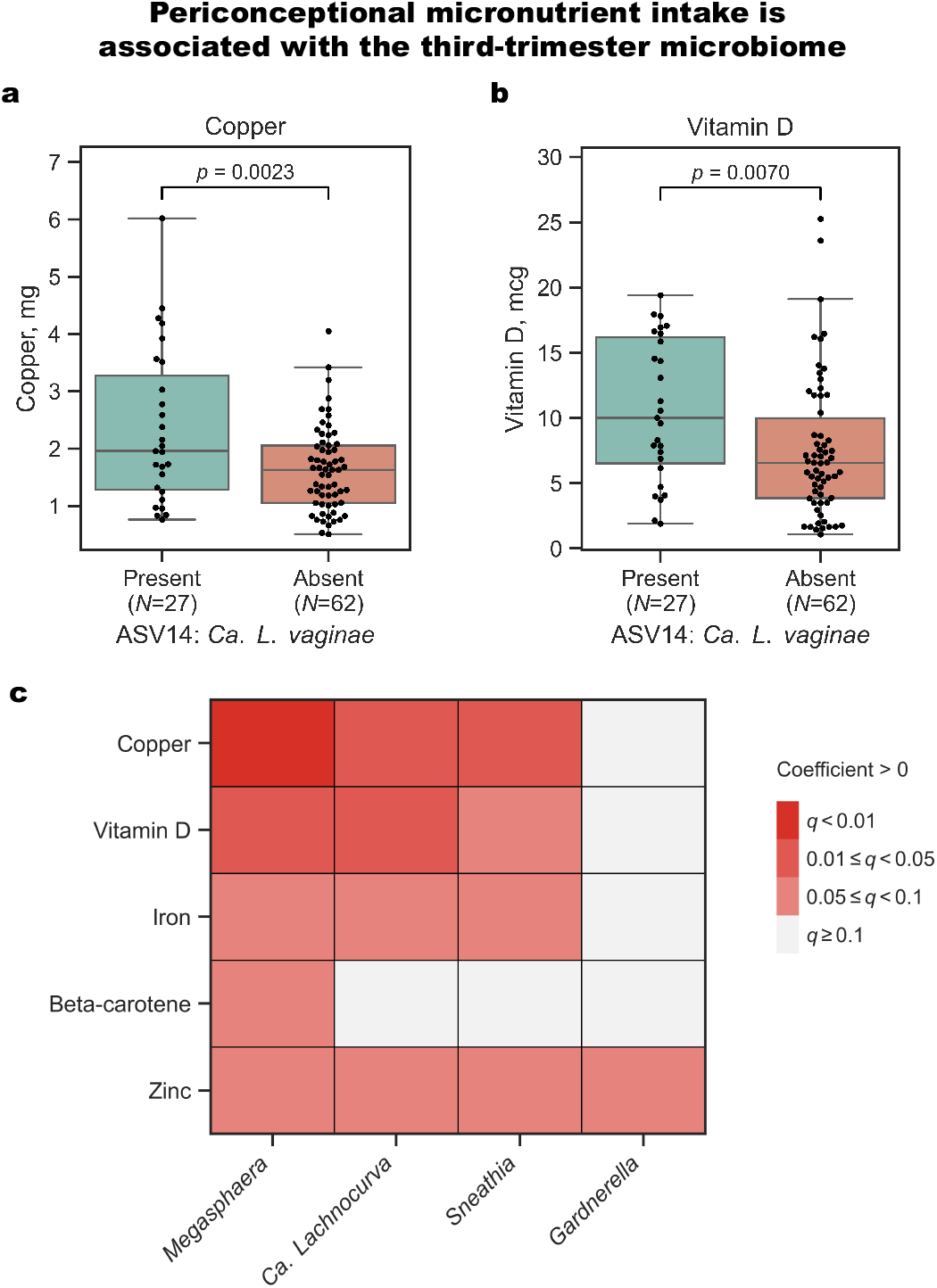
Periconceptional micronutrient intake is associated with prevalence of vaginal taxa in the third trimester. **a,b.** Box and swarm plots (box, IQR; line, median; whiskers, 1.5xIQR) showing total intake of periconceptional copper (**a**) and vitamin D (**b**) grouped by the presence of ASV14: *Ca. L. vaginae* (binomial regression; FDR < 0.1). Presence and absence are called according to a randomly selected rarefaction run for visualization purposes (**Methods**). **c.** Heatmap showing significance of associations between micronutrients and genus prevalence in visit 3 samples (binomial regression; FDR < 0.1).

**Table 1.**
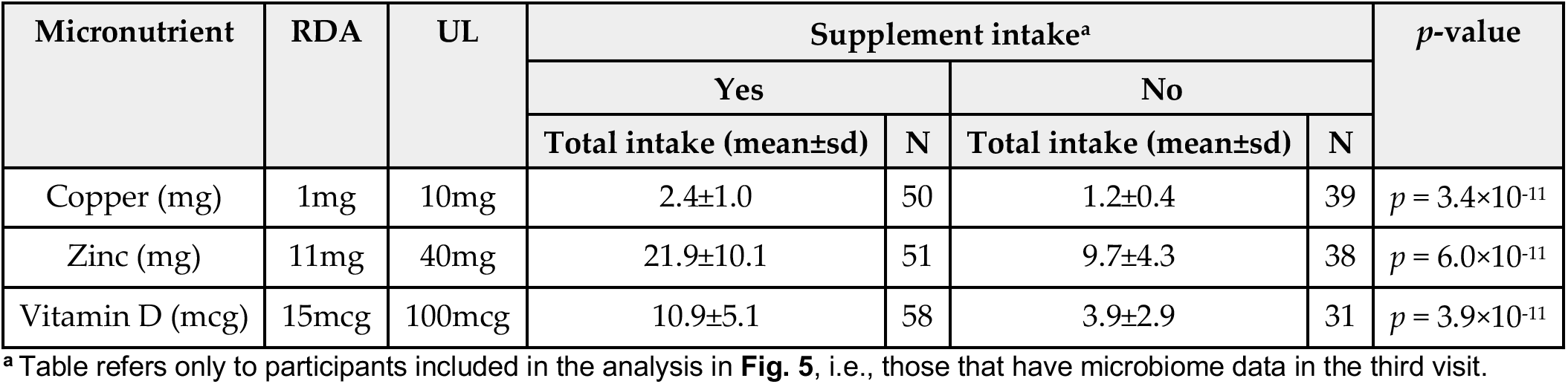
Total daily intake of selected micronutrients stratified by supplement intake.

| **Micronutrient** | **RDA** | **UL** | **Supplement intake<sup>a</sup>** |  |  |  | ***p*-value** |
|---|---|---|---|---|---|---|---|
|  |  |  | **Yes** |  | **No** |  |  |
|  |  |  | **Total intake (mean±sd)** | **N** | **Total intake (mean±sd)** | **N** |  |
| Copper (mg) | 1mg | 10mg | 2.4±1.0 | 50 | 1.2±0.4 | 39 | *p* = 3.4×10<sup>-11</sup> |
| Zinc (mg) | 11mg | 40mg | 21.9±10.1 | 51 | 9.7±4.3 | 38 | *p* = 6.0×10<sup>-11</sup> |
| Vitamin D (mcg) | 15mcg | 100mcg | 10.9±5.1 | 58 | 3.9±2.9 | 31 | *p* = 3.9×10<sup>-11</sup> |

<sup>**a**</sup> Table refers only to participants included in the analysis in **Fig. 5**, i.e., those that have microbiome data in the third visit.

### Microbiome associations with diet and supplement remain in individuals without APOs

To assess whether the overrepresentation of APOs (**Table S1**) influenced our results, we repeated our analyses in participants without APOs, while noting a substantial decrease in our statistical power (N = 48 participants; **Methods**; **Fig. S13a**). First, we evaluated the associations we identified between HEI-2010 scores and the presence of ASV38 (*F. magna H*) and *Corynebacterium* (**Fig. 2**), as ASV65 (*P. harei*) had low prevalence in the 48 participants without APOs. We found that *Corynebacterium* remained significantly associated with HEI-2010 (binomial regression *p* = 0.0058; linear mixed model *p* = 0.020), but not ASV38 (binomial regression *p* = 0.19, linear mixed model *p* = 0.24; **Fig. S13b,c**). Next, we evaluated the associations we identified between dietary components and the first-trimester vaginal microbiome (**Fig. 3a**). We found that 100% fruit juice intake remained significantly associated with microbiome composition (PERMANOVA of Bray-Curtis dissimilarities *p* = 0.037), whereas the associations of niacin and tomatoes were no longer statistically significant (*p* = 0.10 and *p* = 0.059, respectively; **Fig. S13d**). Similarly, our analyses evaluating associations between these components and specific taxa (**Fig. 3b-d**) did not find any significant associations in this smaller subset of participants (ALDEx3; *p* > 0.05 for all; **Fig. S13e-g**). Nonetheless, among the seven supplement groups significantly associated with the third-trimester microbiome (**Fig. 4a**), we observed that six remained significant in the subset of participants without APOs (PERMANOVA of Jaccard distances; *p* < 0.05; **Fig. S13h**). Once again evaluating associations of supplement clusters with the presence of specific taxa (**Figs. 4c,d, S13i**), we found that while the direction of association was the same for all seven associations, only the enrichment of ASV12 (*L. mulieris*) in cluster III remained significant in participants without APOs (*p* = 0.039; *p* > 0.05 for all others; **Fig. S13j,k**). Lastly, we tested the positive associations between total micronutrient intake and the presence of dysbiosis-associated taxa in the third-trimester microbiome (**Fig. 5**), and found that most associations remained statistically significant, including those between ASV14 (*Ca. L. vaginae*) in the third trimester, copper, and vitamin D (binomial regression *p* = 0.012 and 0.0023, respectively; **Fig. S13l,m**), as well as several associations with *Megasphera*, *Ca. Lachnocurva*, and *Sneathia* (**Fig. S13n**). Overall, while bigger population-level studies are needed to further evaluate the associations we found, this analysis suggests that the enrichment of APOs within this pilot cohort did not substantially impact our results.

## Discussion

In this study, by leveraging FFQs targeting the periconceptional period and longitudinal vaginal 16S rRNA amplicon sequencing along the pregnancy of 101 individuals, we investigated the associations between periconceptional nutrition and the vaginal microbiome during pregnancy. We detected associations between dietary indices and specific vaginal microbes only in the first trimester. We further identified fruit juice, tomatoes, and niacin from the diet to be associated with ASVs corresponding to two predominant vaginal *Lactobacillus* species, *L. crispatus* and *L. iners*. For each of these dietary components, the associations with the two *Lactobacillus* species were in opposite directions. Lastly, we found that, in the third trimester, dysbiosis-associated taxa were more likely to be present in individuals with high micronutrient supplementation, and that this correlation was related to copper, zinc, and vitamin D consumption.

Although dietary indices have been widely used for their convenience in summarizing overall diet quality, we only found limited associations between dietary indices and specific vaginal taxa in the first trimester. Notably, several taxa that were positively associated with dietary indices were dysbiosis-associated^52,53^. Together, the limited associations with vaginal microbiome composition and the unexpected direction of taxon-level associations indicate that those indices may not be optimal for capturing factors in diet that are important for the vaginal microbiome during pregnancy. Further, as these associations were primarily at the first trimester, and as vaginal dysbiosis in early pregnancy is associated with spontaneous preterm birth^9^, our findings could indicate a complex interaction between periconceptional diet, APO risk, and the vaginal ecosystem. However, associations of the vaginal microbiome with both dietary indices and specific dietary components were no longer found in later trimesters, indicating that the interplay between diet and vaginal microbes could be transient, while other previously identified endogenous (e.g., hormonal change, stress condition) and exogenous (e.g., exposure to xenobiotics) factors may have stronger effects^2,7,54^.

We found several dietary factors to be differentially associated with the abundance of ASVs corresponding to *L. crispatus* and *L. iners*, with many of these associations occurring in opposite directions. The opposing associations are noteworthy because, despite belonging to the same genus, the two species have distinct ecological characteristics. *L. crispatus* is often considered a marker of a stable, healthy community^52,55^, whereas *L. iners*-dominant communities are associated with elevated risk of transitioning to vaginal dysbiosis^56–59^. The differences in observed associations may be attributable to nutrient utilization and response to the vaginal environment, which differ between the two species^60,61^. We found that niacin exhibited opposite associations with *L. crispatus* and *L. iners*, supporting previous *in vitro* findings demonstrating host dependency of *L. crispatus* on niacin and pointing to distinct degrees of nutritional dependencies between the two species on the host^62^. We also found that intake of tomatoes, the primary source of the antioxidant compound lycopene, was positively associated with *L. crispatus*^63^. This is consistent with previous findings that antioxidant properties in Mediterranean diets are linked to decreased odds of vaginal dysbiosis and APOs^22^, although we did not find evidence of a strong association between aMED scores and the vaginal microbiome. Collectively, these observations suggest that dietary components may contribute to differences in the growth of *Lactobacillus* species, although further characterization and validation are warranted.

While prenatal supplementation has been widely acknowledged as beneficial for maternal and fetal health^64–66^, we found that high intake of periconceptional supplements, such as copper, zinc, and vitamin D, is associated with a higher prevalence of dysbiosis-associated taxa, including *Ca. L. vaginae*, *M. lornae*, *Sneathia*, and *Gardnerella*. This was particularly noteworthy for copper and zinc, in which we found consumption over the RDA, consistent with findings from another US study^67^. The RDA establishes a minimum recommended level of intake to meet the needs of micronutrients in healthy individuals^46,66^. It is suggested that individuals take supplements during pregnancy to reach a total intake substantially higher than the RDA while not exceeding the tolerable upper intake level to optimize reproductive outcomes^66,68^. Therefore, it is surprising to observe that copper and zinc intake at approximately twice their respective RDA, but substantially lower than their tolerable upper intake levels, were already associated with dysbiosis-associated taxa in the vagina. A recent study found that excess maternal circulating copper concentrations during early- and mid-pregnancy are positively associated with the risk of spontaneous preterm birth, potentially reflecting heightened inflammation^69^. Consistent with its potential link to inflammation, we observed taxa associated with vaginal inflammation, including *Sneathia* and *Megasphaera*^70,71^, were enriched in individuals with high copper intake. Additionally, another recent preprint reported that the host cell response to copper and zinc is negatively associated with the abundance of dysbiosis-associated taxa, particularly *Megasphera* and *Gardnerella*, whereas host stress response is positively associated with these taxa, revealing connections among trace metal concentration, host stress response, and dysbiosis of the vaginal microbiome^72^. While this could lead to interesting hypotheses, such that interactions between host and dysbiosis-associated taxa may be related to the host’s response to excess nutrient intake, oxidative stress, and subsequent APO, we caution that, to our knowledge, there is currently no clear evidence that oral intake of these micronutrients is correlated with their vaginal levels^73–75^. Our results suggest that deeper investigation of the full downstream effect of dietary supplementation during pregnancy may be warranted.

Our study has several limitations. First, the incidence rates of APOs in our cohort are significantly higher than the ones observed in the general population^76^, potentially limiting the generalizability of our findings to healthy pregnant individuals. While many of the associations we found remained statistically significant within a subset of participants without APOs, it is difficult to say if others did not replicate because they are influenced by APOs or because of the limited sample size. Second, the dietary and supplement information used in the analysis above is self-reported and collected retrospectively, and may therefore contain minor inaccuracies and be subject to recall biases, despite reflecting general dietary patterns. We note that while such noise in data collection may hide or weaken associations, it is unlikely to cause spurious ones. Third, due to the unavailability of longitudinal dietary profiles, we were unable to account for shifts in dietary profile after the first trimester visit. Finally, we only have information on the total amount of each micronutrient consumed from supplements, and are blind to detailed intake reasons or indications, which might confound the analysis. Collectively, future studies should incorporate high-resolution longitudinal nutritional profiles paired with medically confirmed reasons for supplementation to improve the robustness of similar findings.

In summary, these results demonstrated that periconceptional supplements and certain dietary components are associated with key vaginal microbial taxa at different stages of pregnancy. Future studies integrating longitudinal dietary assessment, metabolomics, hormonal profiling, and strain-level microbiome characterization may help clarify the biological mechanisms underlying these associations and potentially lead to clinically useful nutritional modification strategies.

## Methods

### nuMoM2b cohort

The vaginal samples included in this analysis were collected and banked as part of the nuMoM2b cohort^40^. Between 2010 and 2013, the nuMoM2b study enrolled 10,038 nulliparous pregnant women at 6-14 weeks’ gestation from 8 US clinical sites. Pregnancy dating was confirmed through ultrasound at the enrollment visit. Participants were then followed through delivery. At each of the three study visits during pregnancy (6^0^-13^6^, 16^0^-21^6^, and 22^0^-29^6^ weeks^days^ of gestation), a vaginal swab was self-collected and subsequently suspended in 1 mL 10 mM Tris-HCl, 0.1 mM EDTA. This current analysis was approved by the IRB of Columbia University, approval number AAAT7071.

### Dietary data processing

At study visit 1, dietary intake for the 3 calendar months before pregnancy was assessed using a modified self-administered Block 2005 food frequency questionnaire (FFQ), available in English and Spanish. The FFQ assessed 52 nutrients and 35 food groups from approximately 120 food and beverage items, as well as regular use of dietary supplements. NutritionQuest (Berkeley, CA) performed the subsequent scanning and nutrient mapping for the completed FFQs. In addition, the aMED, HEI-2010, AHEI-2010 total scores were made available for each participant. At each visit, as part of a separate assessment capturing medication use, prenatal vitamin use during pregnancy was recorded qualitatively. These responses were then processed and used to determine the start time of use for each participant^77^.

We filtered out participants with abnormal total intake (<500 or >4000 kcal per day)^16,78^. Then, for micronutrients that can be taken from both diet and supplements, the total intake was calculated by adding up the amount from both sources. Each nutrient or food category was then normalized per 1000 kcal dietary intake per day. Food or nutrients not taken by more than 80 individuals were excluded from downstream statistical analyses. Two nutrients and nine food groups were also excluded because of overlap with other variables in the same category, and additional nine nutrients and six food groups were excluded because they were a-priori considered too fine-grained for downstream analysis, including different types of polyunsaturated fatty acids and specific categories of meat, leaving 59 dietary variables (**Table S2**). Finally, pairwise Pearson’s correlation was calculated between micronutrients taken from dietary supplements, and highly correlated supplements (*r* > 0.95) were grouped. A representative supplement was randomly selected from each group, and its values were used to represent the entire group.

### Clustering of supplement intake profile

To account for differences in scales and units across supplement groups, values within each group were first converted to percentiles. Clustering of supplement groups was done using hierarchical clustering (scipy.cluster.hierarchy 1.10.1) with Euclidean distance and Ward linkage. The number of clusters corresponding to the highest Silhouette score was considered optimal and selected for downstream analysis^79^ (**Fig. S10**). Clusters with fewer than 5 samples were excluded from subsequent analysis, and clusters with 5-10 samples were merged into the cluster with the most similar supplement intake profile. This resulted in four clusters, which were named in ascending order of the mean supplement intake among samples within each cluster (**Table S3**).

### Covariates

At enrollment, demographic data was collected through self-administered questionnaires. In our microbiome analysis, we included the following clinical variables as covariates, due to their potential relevance to the dietary patterns and the vaginal microbiome composition: age at visit 1, education, income as a percentage of the 2013 federal poverty level, and area deprivation index (**Table S1**).

Missing values of income and area deprivation index were imputed using the mean value of the full nuMoM2b cohort. Since our sample size was relatively small, we further collapsed the education level into ‘high school or below’ and ‘over high school’.

### Associations between supplement intake clusters and demographic variables

Associations between supplement clusters and the aforementioned demographic variables were tested using Fisher’s exact tests for categorical variables and Kruskal-Wallis tests for continuous variables.

Fisher’s exact tests were performed using the stats package in R (v4.3.3), accessed through Python package rpy2 (v3.5.11), while Kruskal-Wallis tests were performed using the scipy.stats (v1.10.1) package in Python. False discovery rate (FDR) corrections were done once using the Benjamini-Hochberg method for all tests^80^. Variables showing significant overall associations with supplement clusters were selected for pairwise comparisons between supplement clusters. Pairwise comparisons were performed between cluster I (baseline) and each of the other clusters using Fisher’s exact test for categorical variables and Mann-Whitney U test for continuous variables. FDR corrections were then done per variable.

### 16S rRNA gene sequencing of vaginal swabs

Microbial DNA was extracted from 352 vaginal swabs in 4 batches. The V3-V4 region of the 16S rRNA gene was amplified using forward primer CCTACGGGNGGCWGCAG and reverse primer GACTACHVGGGTATCTAATCC. The resulting libraries were barcoded and sequenced using MiSeq Reagent kit v3 (600-cycles) on the MiSeq platform to a mean±std depth of 28,834±15,009 reads.

All reads were aligned to T2T-CHM13 and PhiX genomes using bowtie2^81^, and mapped reads were removed to limit off-target analysis. Primers were trimmed using cutadapt v5.1 (ref. ^82^). DADA2 v1.30.0 was then used to remove low-quality reads, merge paired-end reads, and count ASVs present in samples^83^. Overall, 2,332 ASVs were identified. Potential contamination contained in the samples was modeled and removed *in silico* for each batch using SCRuB^84^. Seven samples containing processing errors were identified by Metagenome Matcher on corresponding metagenomic data and were removed^41^. Taxonomy was assigned to ASVs for interpretability using speciateIT with the vSpeciateDB V3V4 model^85^. As speciateIT uses a 7th-order Markov Chain model and provides a posterior probability to each taxonomy assigned, taxonomic annotations with posterior probabilities >0.85 were considered as high confidence, and others were subsequently re-annotated as ‘Unknown’. We further filtered out rare ASVs, defined as those not present in at least 3 samples with more than 5 reads. 545 ASVs were retained. Abundance tables at the genus level were generated by collapsing all ASVs with the same genus-level annotation.

Rarefaction of microbial reads was done to support beta-diversity (inter-sample distance) estimation and taxon presence/absence determination. To balance rarefaction depth and sample size, 11 samples with <5000 reads were dropped, and all other samples were rarefied to 5000 reads. To compensate for the instability of taxon presence due to low rarefaction depth, the rarefaction process was repeated 1000 times. For beta-diversity estimates, the Bray-Curtis dissimilarity and Jaccard distance between samples were calculated for each iteration, and the mean distance across all iterations was used as the final estimate. Taxon presence/absence was represented as the proportion of rarefaction iterations in which the taxon was detected.

### Associations between diet and vaginal microbiome

In addition to age at visit 1, education, income as a percentage of the 2013 federal poverty level, and area deprivation index (**Table S1**), as detailed above, we additionally adjusted for diet quality by including AHEI-2010 total score as a covariate when investigating the associations between dietary supplements and vaginal microbiome. Beta-diversity comparisons were conducted via permutational multivariate analysis (PERMANOVA) using the ‘adonis2’ function in the R package vegan v2.7.2. FDR corrections were done using the Benjamini-Hochberg method^80^. FDR corrections were done per visit and per beta-diversity metric, and were separate for diet (including nutrients and food groups) and dietary supplements (**Table S2**).

Differential abundance analyses were performed to assess the associations of dietary indices and components with the relative abundance of vaginal microbial taxa in R 4.3.3 using the Dirichlet-multinomial model from ALDEx3^86^. We assumed no scale differences between samples and applied total sum scaling with gamma=0.5. We included only microbes that were present in at least 25% of samples, and collapsed other taxa into one “others” category, resulting in 16 ASVs and 16 genera. Since ALDEx3 generated near-uniform *p*- and *q*-values of approximately 0.5 when the range of a continuous variable exceeded 10, we divided all continuous variables by the largest power of ten that was smaller than the max value of the variable, forcing the values to be distributed between 0 and 10. We then scaled back coefficient estimates. Presence/absence analyses were performed using the binomial regression model from scipy.stats (v1.10.1). We compared only taxa that were present in between 30- 70% of the samples for each test to reduce the influence of sparse features and increase statistical power.

In linear mixed modeling, as samples were from different pregnancy visits, gestational age was also included as a covariate. We used the same minimum and maximum prevalence thresholds as described above. When testing on the differential abundance of ASVs, we first applied the centered log-ratio transformation to the relative abundance of all ASVs after adding a pseudocount of 5e-5, then ran a linear mixed model from statsmodels.formula.api.mixedlm (v0.14.1). When testing on the presence of taxa, the same binomial regression was applied as in per-visit analyses, except that samples were additionally grouped by participant. All analyses were done in Python 3.8.20.

### Associations restricted to individuals without APOs

To assess whether our findings were affected by the overrepresentation of APO cases in the cohort, we restricted the samples to the ones from individuals without APOs and tested if the main findings in this study remain statistically significant in this subset (**Fig. S13**). Significant associations (FDR < 0.1) in the full cohort were tested again on this subset following the same procedure, with statistical significance in the subset defined as *p* < 0.05. Association between HEI-2010 score and the presence of ASV65 (*P. harei*) could not be retested, because the prevalence of the ASV was below the minimum prevalence threshold (30%) in the subset. We were also unable to retest the associations between micronutrient intake and the presence of *Gardnerella* at visit 3, because its prevalence exceeds the maximum prevalence threshold in the subset (70%).

## Acknowledgments

We thank members of the Korem and Uhlemann groups for useful discussions. We thank William F. Kindschuh for contributions to cohort design and data generation. We thank all participants, staff, and investigators involved in sample collection and generation of data used in this study. This study was supported by grant funding from the Eunice Kennedy Shriver National Institute of Child Health and Human Development (NICHD): R01 HD106017 (T.K.), R01 HD114715 (T.K.), F31 HD115394 (J.A.U.), U10 HD063036, U10 HD063072, U10 HD063047, U10 HD063037, U10 HD063041, U10 HD063020, U10

HD063046, U10 HD063048, and U10 HD063053; the National Library of Medicine: T15 LM007079 (J.A.U.); and the Clinical and Translational Science Institutes: UL1 TR001108 and UL1 TR000153.

## Author contributions

J.L. and T.K. conceived and designed the study. J.L. analyzed data, with assistance from A.B.K and J.A.U. H.P. and E.W. prepared and sequenced the microbial samples under supervision by A.-C.U. A.B.K. and J.A.U. processed sequencing data. J.L. and T.K. wrote the manuscript with critical input from all authors. T.K. supervised the study.

## Supplementary figures

**Supplementary Figure 1.**
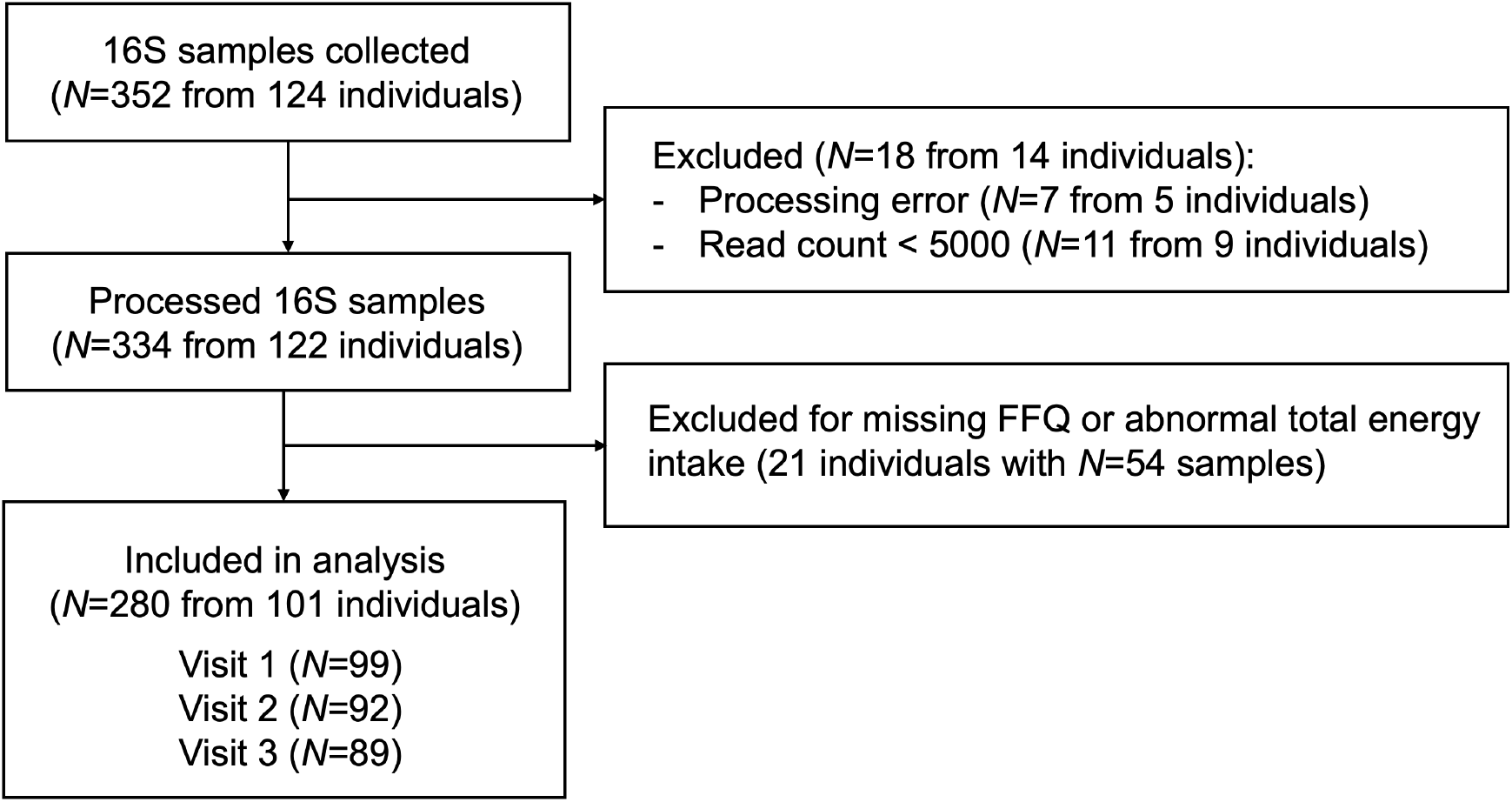
Sample selection workflow. Flow chart of sample exclusion process.

**Supplementary Figure 2.**
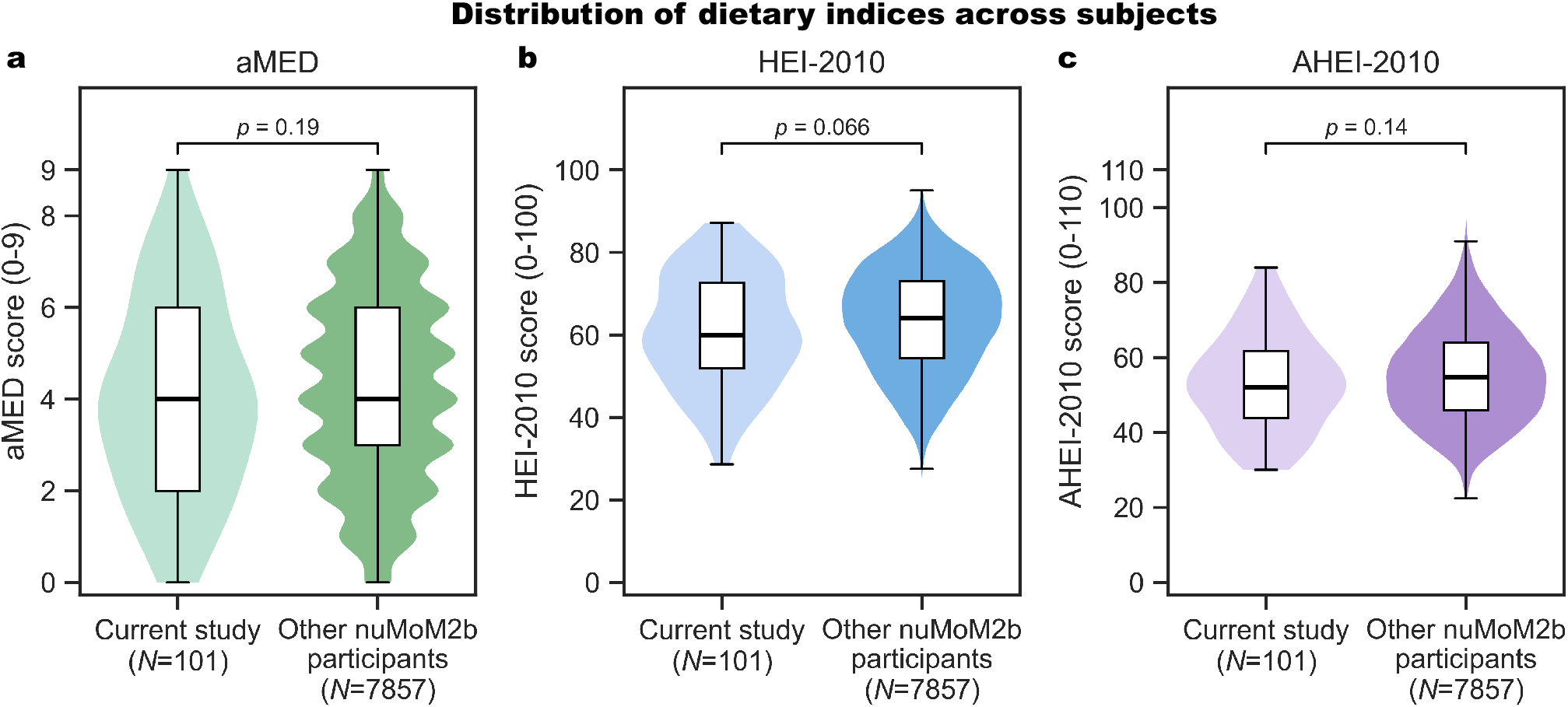
Distributions of dietary indices in the current study are not significantly different from those of other participants in the nuMoM2b cohort. **a-c.** Violin and box plots (box, IQR; line, median; whiskers, 1.5xIQR) of aMED (**a**), HEI-2010 (**b**), and AHEI-2010 (**c**) scores grouped by inclusion status in the current study (two-sided Mann-Whitney U test).

**Supplementary Figure 3.**
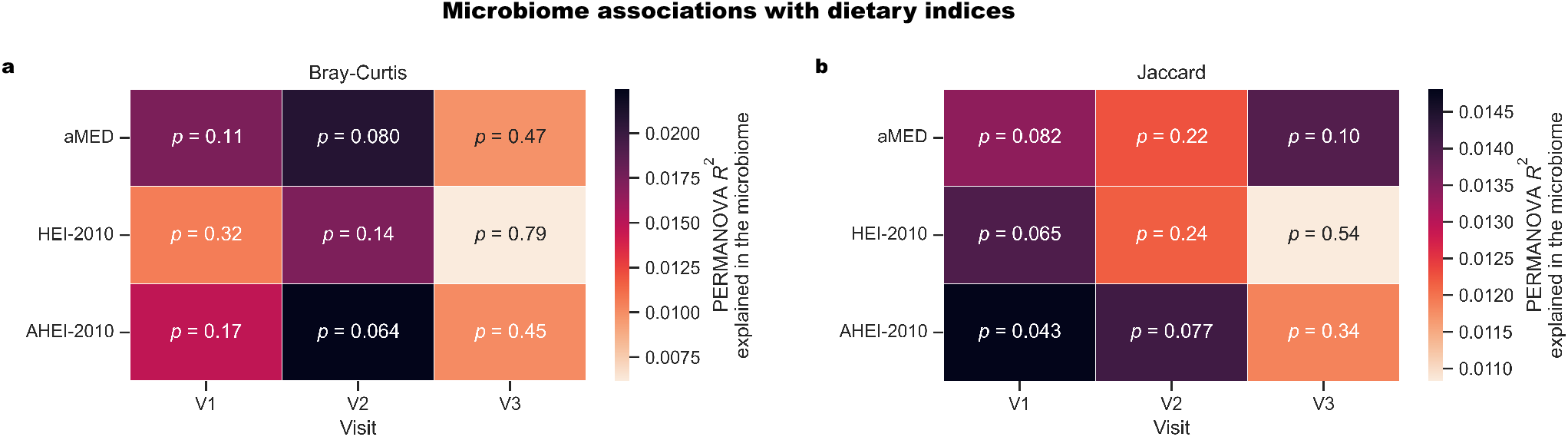
Periconceptional dietary indices are not associated with the composition of the vaginal microbiome during pregnancy. **a,b.** Heatmap of PERMANOVA *R*^2^ between dietary indices and beta-diversity of microbiome across visits, measured using Bray-Curtis dissimilarity (**a**) and Jaccard distance (**b**). PERMANOVA *p* values are listed in each box. FDR > 0.1 for all.

**Supplementary Figure 4.**
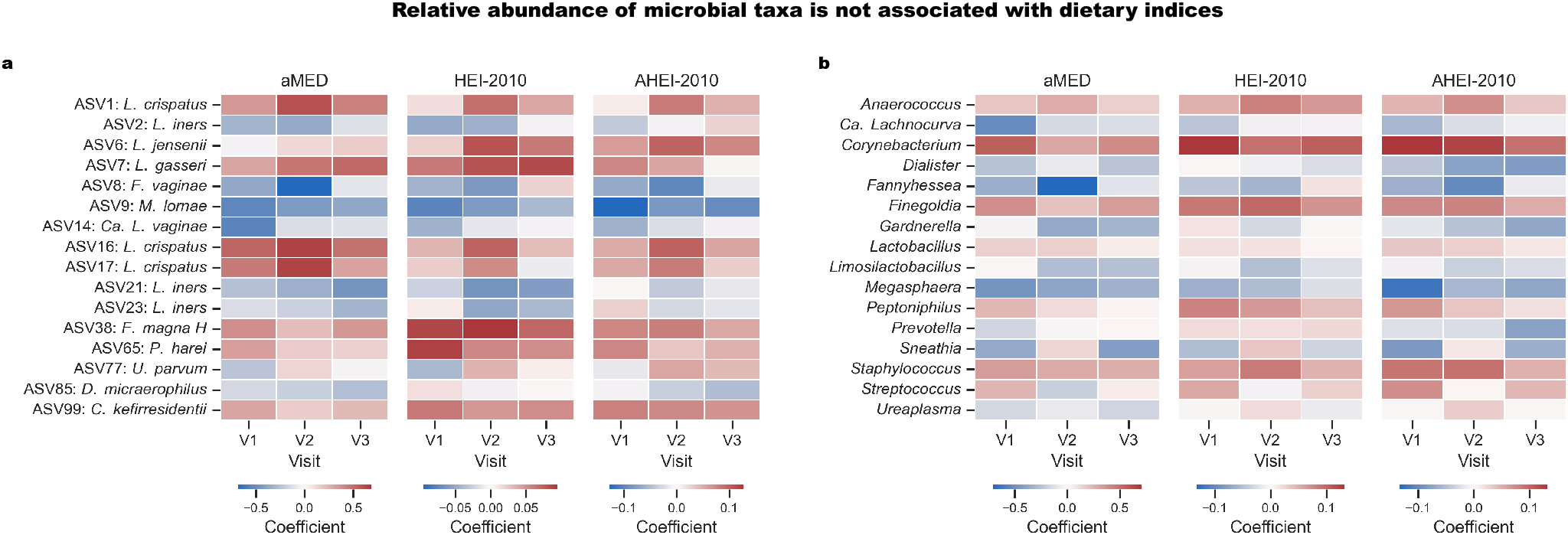
Periconceptional dietary indices are not associated with the relative abundance of vaginal microbes during pregnancy. **a,b.** Heatmaps showing ALDEx3 coefficients between dietary indices and the relative abundance of vaginal microbial ASVs **(a)** and genera **(b)** across samples collected in the three study visits. FDR > 0.1 for all comparisons.

**Supplementary Figure 5.**
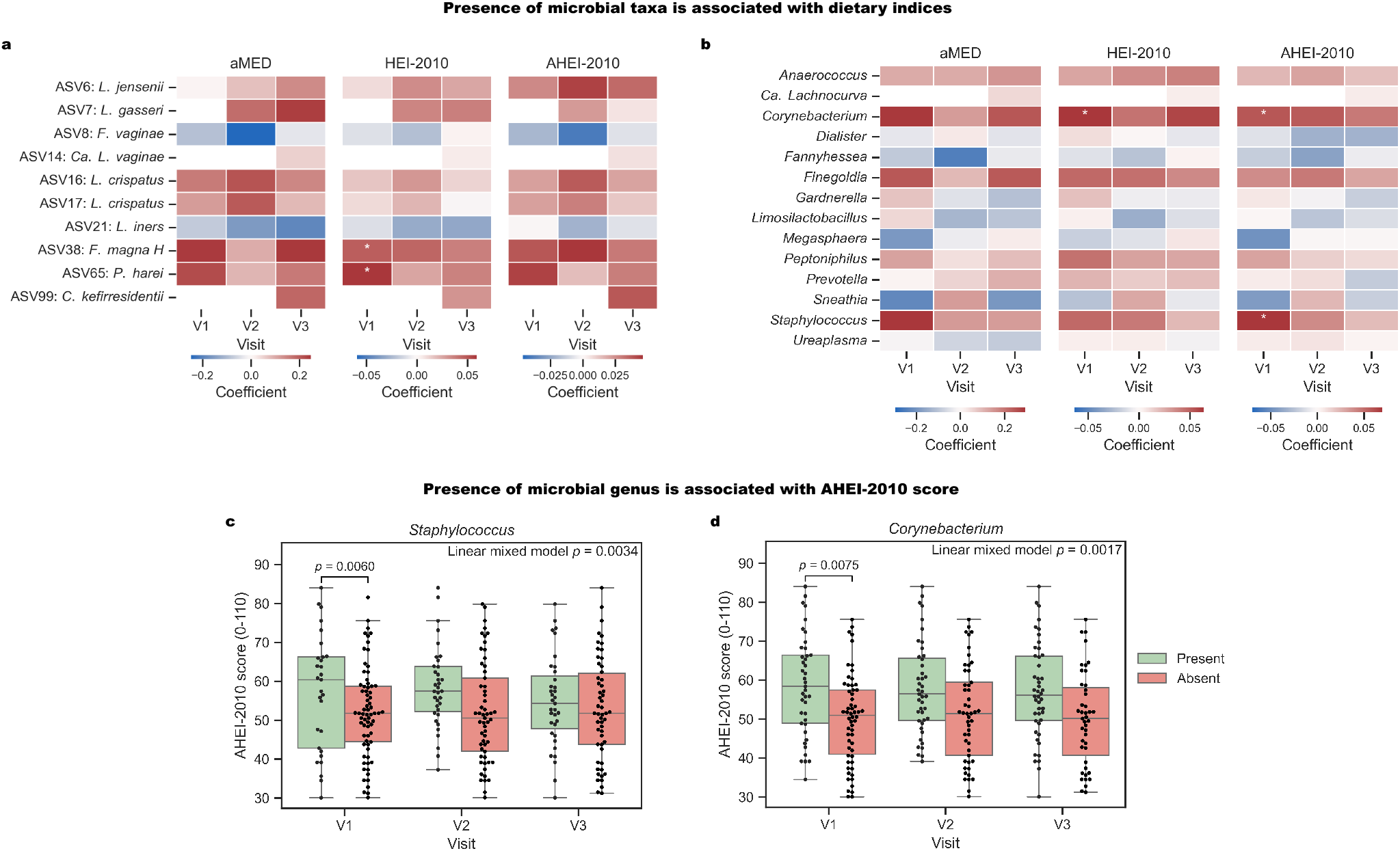
Periconceptional dietary indices are associated with the presence of vaginal microbes in the first trimester. **a,b.** Heatmaps showing binomial regression coefficients between dietary indices and presence of vaginal microbial ASVs (**a**) and genera (**b**) across three visits (*: binomial regression *p* < 0.05; FDR < 0.1). **c,d.** Box and swarm plots (box, IQR; line, median; whiskers, 1.5xIQR) showing AHEI-2010 score grouped by presence of *Staphylococcus* (**c**), and *Corynebacterium* (**d**) across visits. The visit-specific *p*-value is from the corresponding binomial regression model in **b**, while the overall *p*-value of each panel corresponds to a linear mixed model fitted on samples across all visits (FDR < 0.1). While both presence and absence are called according to a randomly selected rarefaction run for visualization purposes, statistical analyses were performed on fraction of presence across rarefaction runs (**Methods**).

**Supplementary Figure 6.**
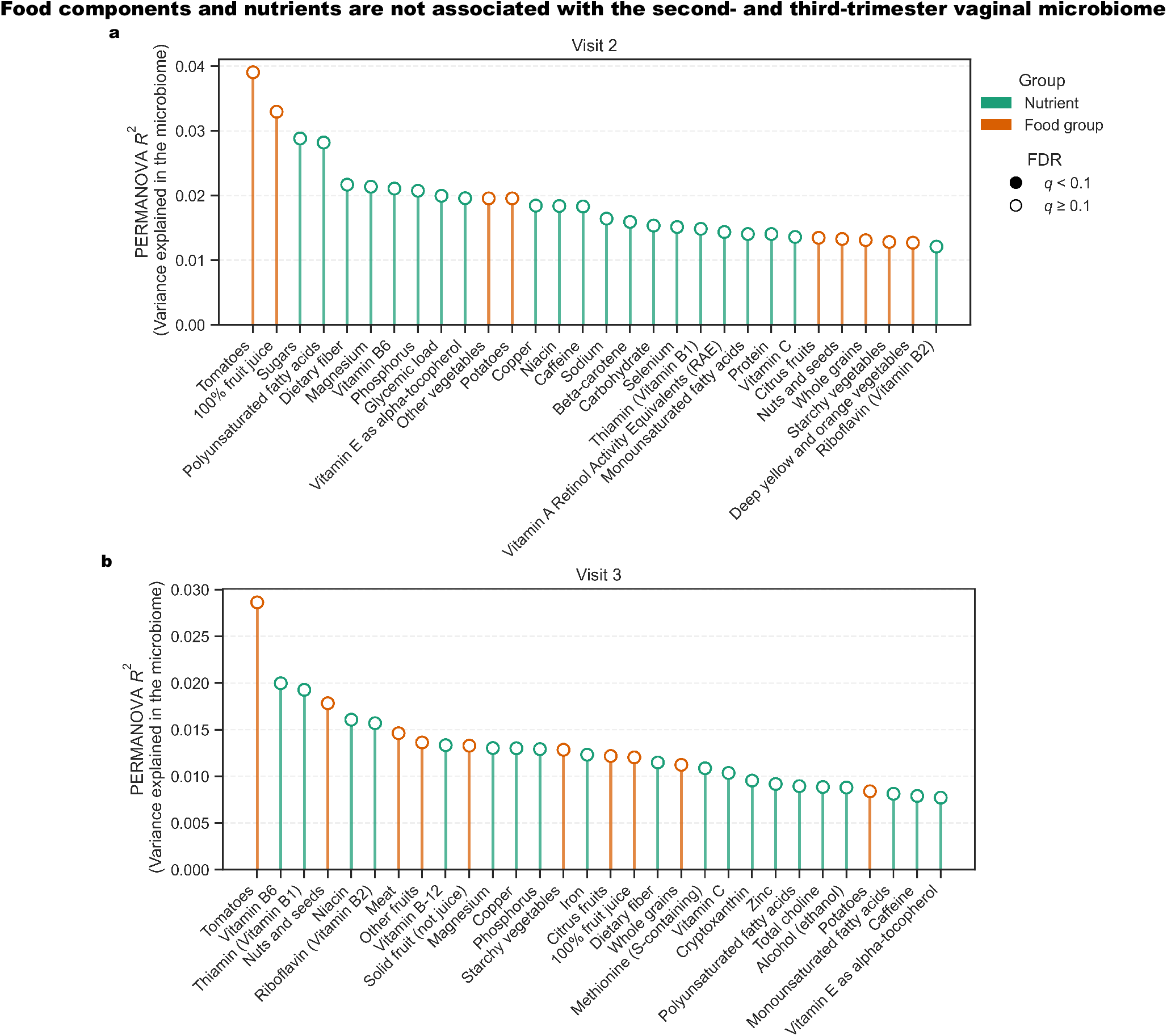
Dietary components are not associated with the second and third trimester vaginal microbiome. **a,b**. Lollipop plot of PERMANOVA *R*^2^ between diet variables (top 30 based on *R*^2^) and Bray-Curtis dissimilarity at visit 2 (**a**) and 3 (**b**).

**Supplementary Figure 7.**
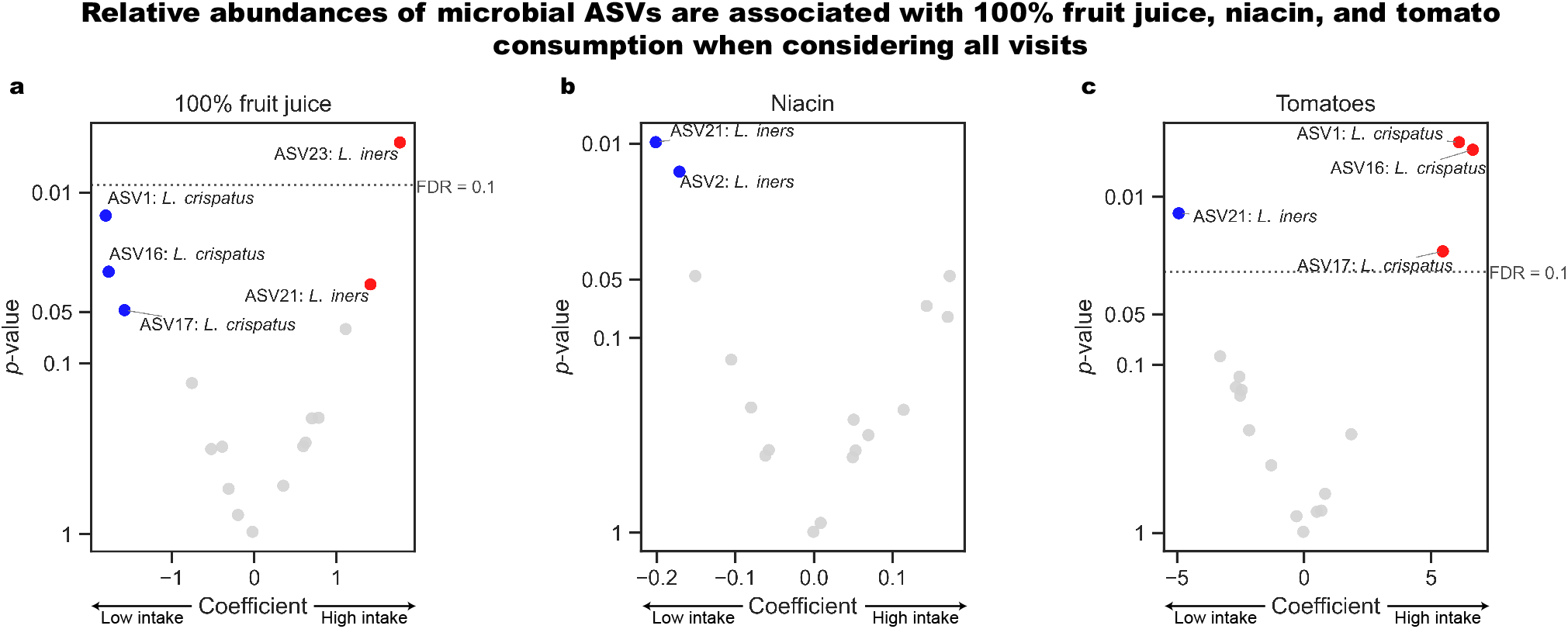
Specific ASVs associated with 100% fruit juice, niacin, and tomato consumption across visits. **a-c.** Volcano plot showing statistical significance (y-axis) and association coefficient (linear mixed model; x-axis) between abundance of microbial ASVs and 100% juices (**a**), niacin (**b**), and tomatoes (**c**).

**Supplementary Figure 8.**
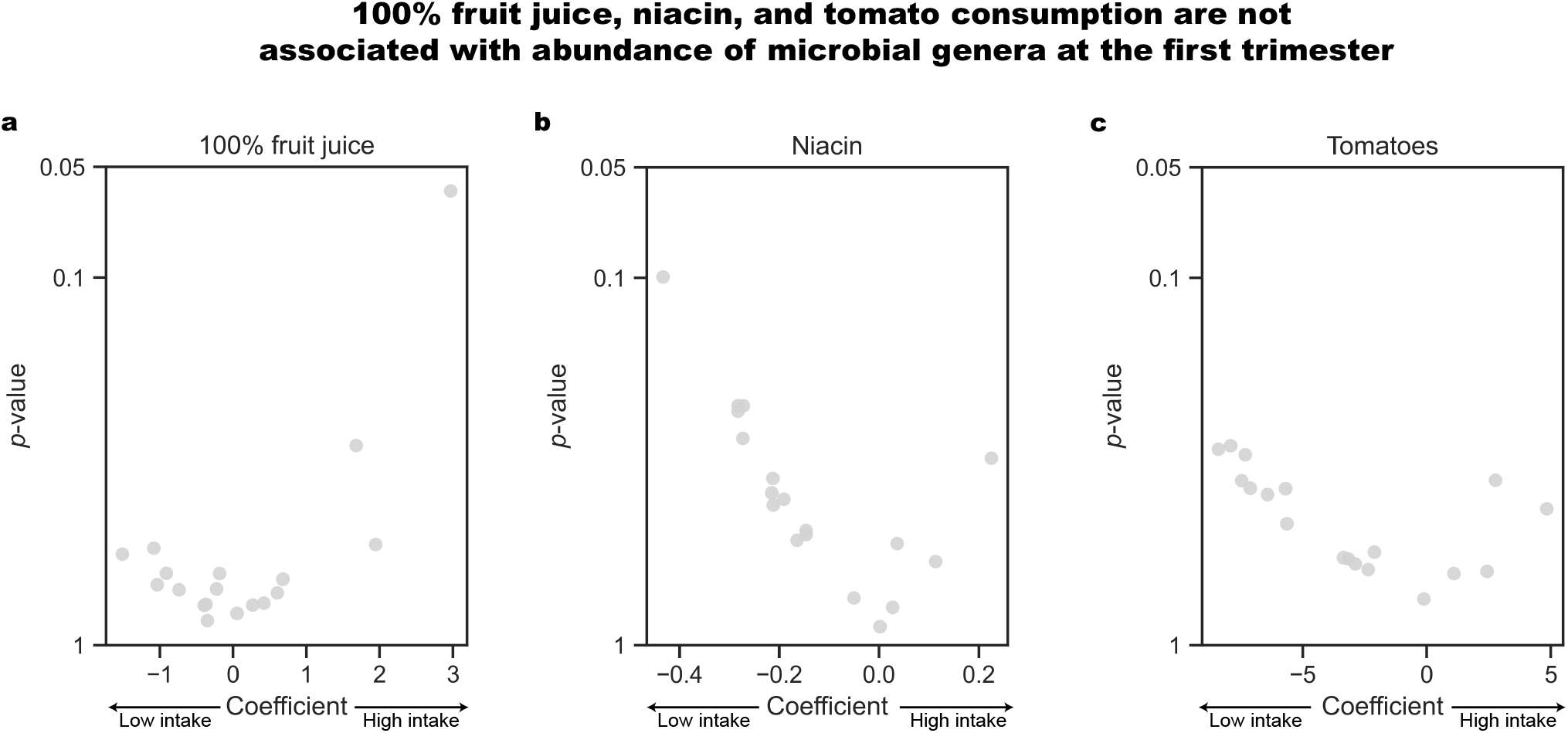
Juice (100% fruit), niacin, and tomato consumption is not associated with abundances of microbial genera in the first trimester. **a-c.** Volcano plot showing statistical significance (ALDEx3, y-axis) and association coefficients (x-axis) between abundance of microbial genus at visit 1 and 100% fruit juices (**a**), niacin (**b**), and tomatoes (**c**). FDR > 0.1 for all.

**Supplementary Figure 9.**
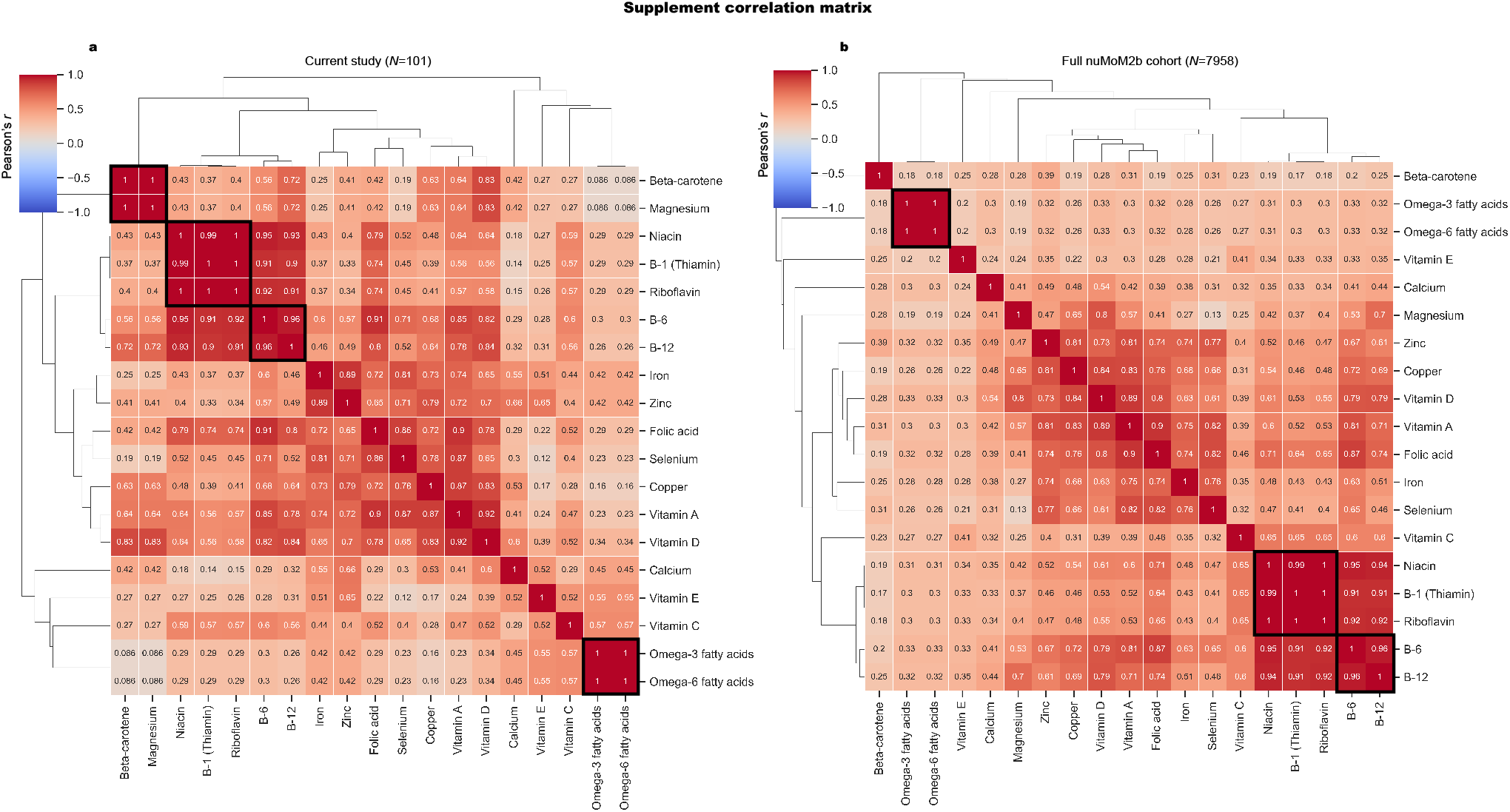
Dietary supplements are highly correlated. **a,b,** Pearson’s correlation matrix between dietary supplements in participants included in this study (**a**) and the full nuMoM2b cohort (**b**). Supplements are hierarchically clustered by Pearson’s correlation using average linkage. Black boxes highlighted along the diagonal correspond to highly correlated supplements (Pearson’s *r* > 0.95) that are subsequently grouped together. Most supplement groups with nearly perfect correlations in this study (**a**) also exhibited similar relationships in the full nuMoM cohort (**b**), except for beta-carotene and magnesium.

**Supplementary Figure 10.**
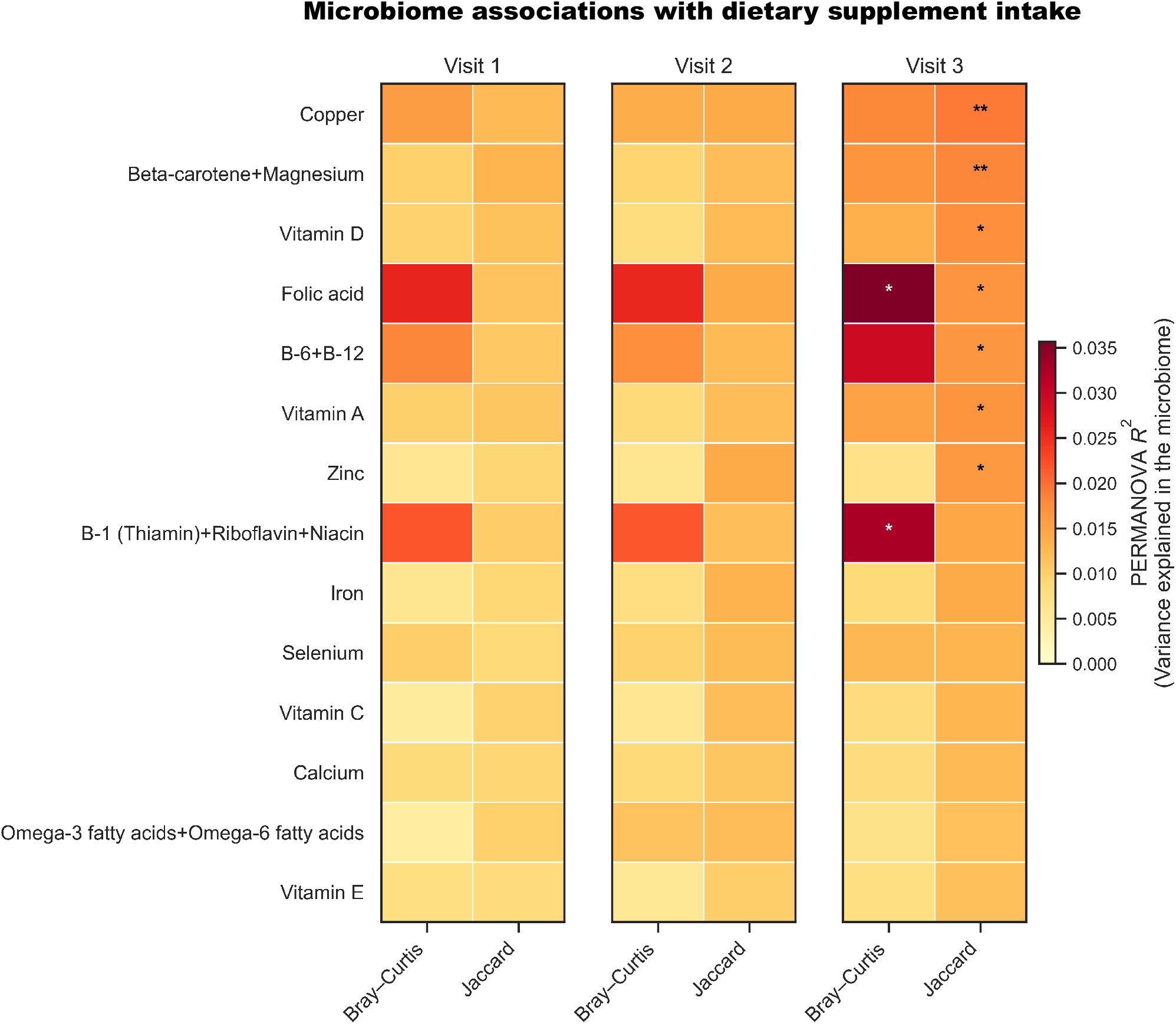
Dietary supplement intake is associated with the vaginal microbiome in the third trimester. Heatmap of PERMANOVA *R*^2^ between diet variables and beta diversity of the microbiome across visits (***: *p* < 0.001, **: *p* < 0.01, *: *p* < 0.05; FDR < 0.1).

**Supplementary Figure 11.**
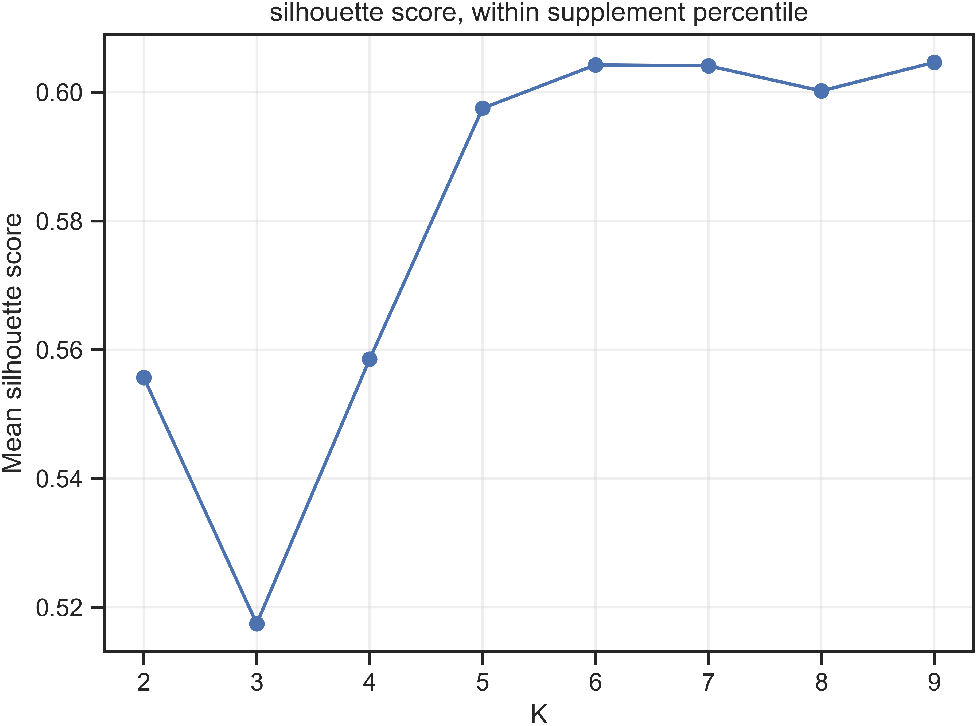
Selection of the optimal number of clusters. Mean silhouette score from hierarchical clustering using Euclidean distance and Ward linkage with number of clusters (K) from 2 to 9.

**Supplementary Figure 12.**
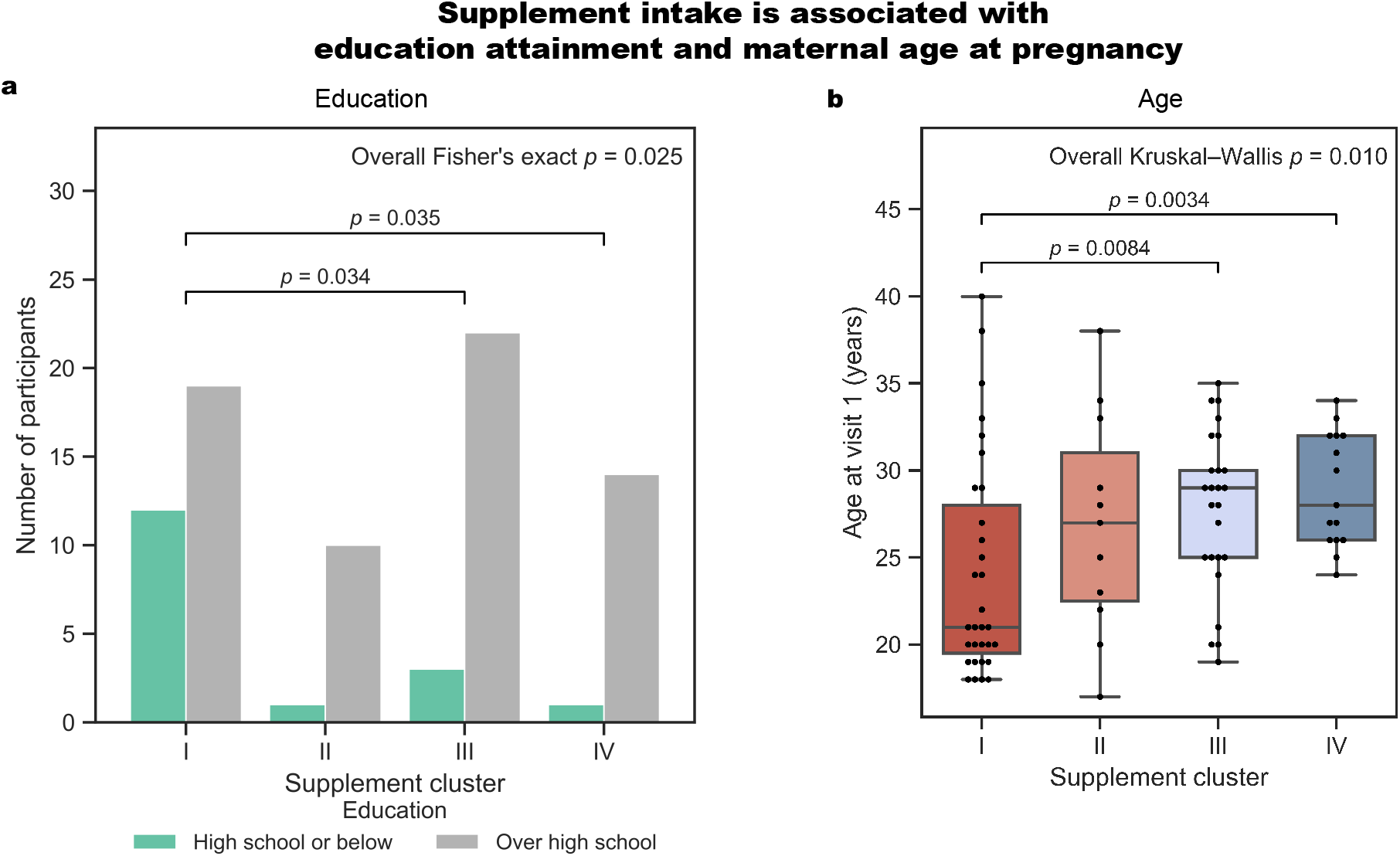
Dietary supplement intake is associated with age and education. **a.** Bar plots of the number of individuals in each supplement cluster, separated by education level. Shown are Fisher’s exact *p* values (FDR < 0.1). **b.** Box and swarm plots (box, IQR; line, median; whiskers, 1.5xIQR) showing maternal age at visit 1, grouped by supplement cluster. The *p* values for pairwise comparisons are from the two-sided Mann-Whitney U test (FDR < 0.1).

**Supplementary Figure 13.**
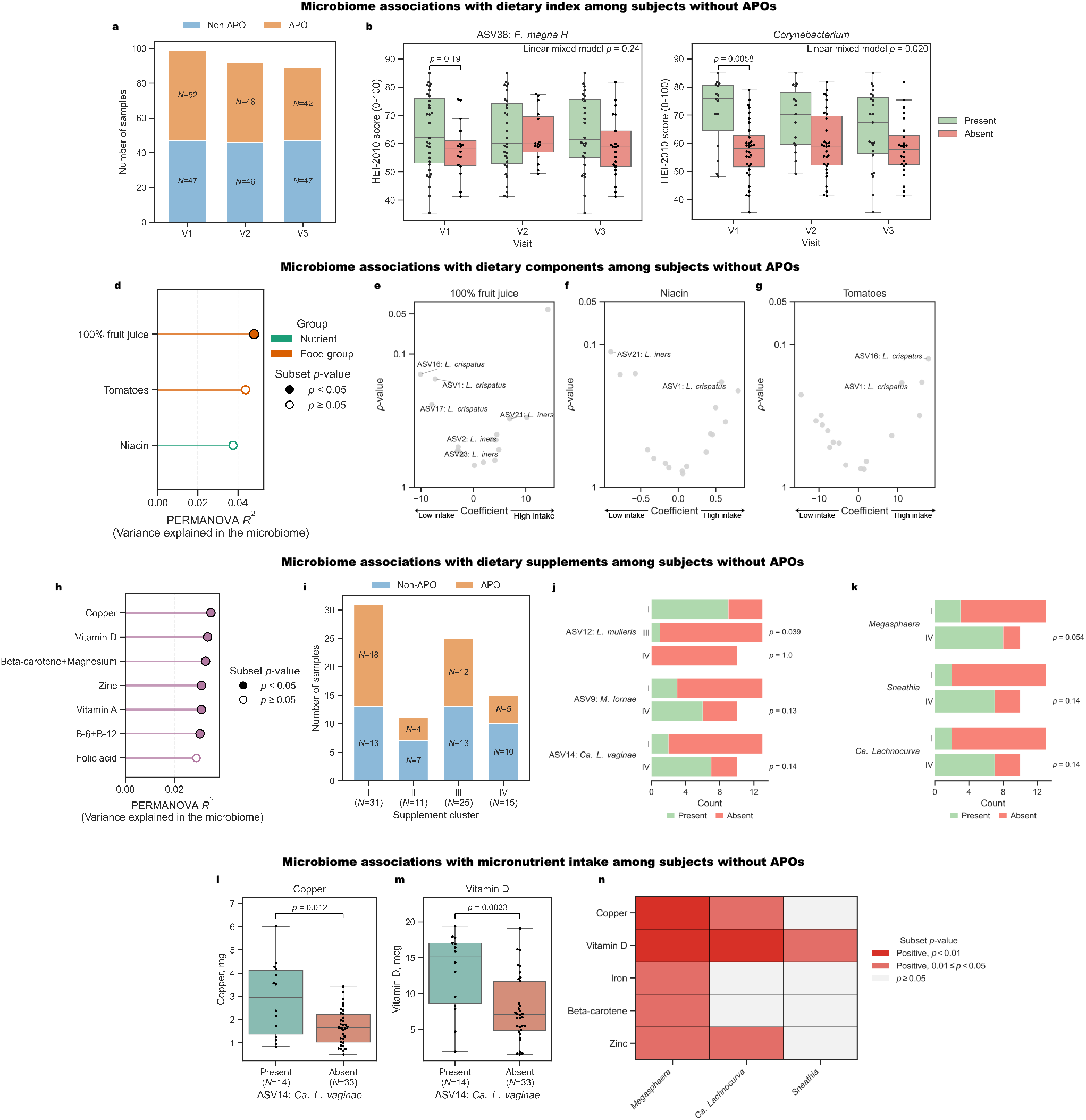
Microbiome associations with dietary patterns remain in participants without APOs. **a.** Stacked bar plot showing distribution of samples with and without APOs across visits. b-h. Evaluation of the analysis in **Figs. 2e,f (b,c), 3a (d), 3b-d (e-g)**, and **4a** (h), restricted to 48 individuals without APOs. i. Stacked bar plot showing distribution of samples with and without APOs across the supplement clusters identified in **Fig. 4b. j-n**. Replication of **Figs. 4c,d (j,k), 5a,b (l,m)**, and **5c (n)**, restricted to individuals without APOs.

## Supplementary tables

**Table S1.** Distribution of sociodemographic variables associated with diet quality and vaginal microbiome.

| **Variable** |  |
|---|---|
| **Age at visit 1 (years, mean±sd)** | 26.45±5.57 |
| **Household Income as Percentage of Federal Poverty Rate (%, mean±sd)** | 434.11±271.25 |
| **Area deprivation index (mean±sd)** | 46.02±27.95 |
| **Education (N [%])** |  |
| High school or below | 79 (78%) |
| Over high school | 22 (22%) |
| **Self-identified Race (N [%])** |  |
| White | 65 (64%) |
| Black | 12 (12%) |
| Others | 24 (24%) |
| **Adverse pregnancy outcomes (APOs; N [%])** |  |
| No APO | 48 (48%) |
| Any APO | 53 (52%) |
| Spontaneous preterm birth | 19 (19%) |
| Fetal demise | 6 (6%) |
| Chorioamnionitis | 1 (1%) |
| Preeclampsia | 20 (20%) |
| Gestational diabetes | 5 (5%) |
| Hypertension | 8 (8%) |
| Small for gestational age | 6 (6%) |
| **Start time of prenatal vitamin intake (N [%])** |  |
| >2 Months before pregnancy | 24 (24%) |
| During the 2 months before pregnancy | 13 (13%) |
| During pregnancy, before first visit | 56 (55%) |
| During pregnancy, after first visit | 3 (3%) |
| Missing or never taken | 5 (5%) |

**Table S2.** FFQ variables included in this analysis.

| **Category** | **Variables** |
|---|---|
| **Nutrients** | Saturated fat; Monounsaturated fatty acids; Polyunsaturated fatty acids; Cholesterol; Glycemic load; Protein; Carbohydrate; Total fat; Alcohol (ethanol); Sugars; Dietary fiber; Total choline; Calcium; Iron; Magnesium; Phosphorus; Potassium; Sodium; Zinc; Copper; Selenium; Methionine (S-containing); Caffeine; Theobromine; Average daily dietary folate equivalents; Vitamin D; Vitamin A Retinol Activity Equivalents (RAE); Retinol; Alpha-carotene; Beta-carotene; Cryptoxanthin; Lycopene; Lutein-Zeaxanthin; Vitamin E as alpha-tocopherol; Vitamin C; Thiamin (Vitamin B1); Riboflavin (Vitamin B2); Niacin; Vitamin B6; Vitamin B-12; Vitamin K as phylloquinone. |
| **Foods** | Milk; Yogurt; Cheese; Citrus fruits, melons, and berries; Other fruits; Solid fruit (not juice); 100% fruit juice; Whole grains; Non-whole grains; Meat, fish and poultry; Nuts and seeds; Dry beans and peas; Dark-green vegetables; Deep yellow and orange vegetables Potatoes; Starchy vegetables;, Tomatoes; Other vegetables. |
| **Supplements** | Vitamin A; Vitamin C; Vitamin D; Vitamin E; Iron; Calcium; Zinc; Beta-carotene+Magnesium; B-1 (Thiamin)+Riboflavin+Niacin; B-6+B-12; Folic acid; Copper; Selenium; Omega-3 fatty acids+Omega-6 fatty acids. |

**Table S3.** Supplement intake profile of each cluster.

|  | **I** | **II** | **III** | **IV** |
|---|---|---|---|---|
| **Vitamin A** | 0.00 ± 0.00 | 338.41 ± 282.57 | 1515.00 ± 0.00 | 2361.96 ± 747.00 |
| **Vitamin C** | 8.64 ± 30.94 | 21.10 ± 19.77 | 185.00 ± 206.20 | 214.21 ± 274.24 |
| **Vitamin D** | 0.00 ± 0.00 | 1.60 ± 1.01 | 5.46 ± 1.24 | 13.26 ± 4.11 |
| **Vitamin E** | 0.00 ± 0.00 | 2.76 ± 2.78 | 6.89 ± 5.35 | 18.38 ± 13.07 |
| **Iron** | 0.00 ± 0.00 | 5.94 ± 8.75 | 26.92 ± 6.82 | 30.38 ± 12.11 |
| **Calcium** | 9.22 ± 51.32 | 17.82 ± 20.08 | 103.43 ± 158.49 | 375.62 ± 366.44 |
| **Zinc** | 0.00 ± 0.00 | 2.18 ± 2.35 | 11.77 ± 2.99 | 19.62 ± 6.32 |
| **Beta-carotene** | 0.00 ± 0.00 | 116.87 ± 125.23 | 0.00 ± 0.00 | 1070.86 ± 343.40 |
| **B-1 (Thiamin)** | 0.00 ± 0.00 | 0.32 ± 0.26 | 2.09 ± 2.41 | 2.27 ± 0.70 |
| **B-6** | 0.00 ± 0.00 | 0.43 ± 0.35 | 2.24 ± 1.20 | 3.05 ± 0.95 |
| **B-12** | 0.00 ± 0.00 | 0.91 ± 0.56 | 3.42 ± 2.89 | 7.09 ± 1.83 |
| **Folic acid** | 12.90 ± 71.84 | 114.54 ± 116.56 | 718.86 ± 213.12 | 764.57 ± 274.61 |
| **Copper** | 0.00 ± 0.00 | 0.23 ± 0.23 | 1.00 ± 0.00 | 2.30 ± 0.75 |
| **Selenium** | 0.00 ± 0.00 | 8.62 ± 12.99 | 60.00 ± 0.00 | 56.38 ± 27.56 |
| **Riboflavin** | 0.00 ± 0.00 | 0.34 ± 0.26 | 2.09 ± 2.41 | 2.45 ± 0.72 |
| **Magnesium** | 0.00 ± 0.00 | 9.74 ± 10.44 | 0.00 ± 0.00 | 89.24 ± 28.62 |
| **Niacin** | 0.00 ± 0.00 | 4.21 ± 3.34 | 24.86 ± 24.09 | 29.85 ± 9.07 |
| **Omega-3 fatty acids** | 0.00 ± 0.00 | 0.00 ± 0.02 | 0.07 ± 0.16 | 0.07 ± 0.18 |
| **Omega-6 fatty acids** | 0.00 ± 0.00 | 0.00 ± 0.00 | 0.01 ± 0.03 | 0.01 ± 0.04 |

